# Protein structure, a genetic constraint on glycosylation

**DOI:** 10.1101/2024.05.15.594261

**Authors:** Benjamin P. Kellman, Daniel Sandoval, Olga O. Zaytseva, Kelly Brock, Sabyasachi Baboo, Daniela Nachmanson, Edward B. Irvine, Sanne Schoffelen, Erick Armingol, Nathan Mih, Yujie Zhang, Mia Jeffris, Philip Bartels, Thi Nguyen, Amy Tam, Sarah Gasman, Shlomi Ilan, Samuel William Canner, Isaac Shamie, Jolene K. Diedrich, Xiaoning Wang, Esther van Woudenbergh, Meghan Altman, Anthony Aylward, Bokan Bao, Andrea Castro, James Sorrentino, Austin W.T. Chiang, Matt Campbell, Yannic Bartsch, Patricia Aguilar-Calvo, Christina Sigurdson, Galit Alter, Gordan Lauc, John R. Yates, Bjørn Gunnar Rude Voldborg, Debora Marks, Frederique Lisacek, Nathan E. Lewis

## Abstract

Unlike DNA, RNA, and protein biosynthesis, dogma describes glycosylation as primarily determined by intrinsic cellular limitations, such as glycosyltransferase expression and precursor availability. However, this cannot explain the commonly-observed differences between glycans on the same protein. By examining site-specific glycosylation on diverse human proteins, we detected associations between protein structure and glycan structure, broadly generalizable to human-expressed glycoproteins. Through structural analysis of site-specific glycosylation data, we found protein-sequence and structural features consistently correlated with specific glycan features. To quantify these relationships, we present a new amino acid substitution matrix describing “glycoimpact”, i.e., the association of primary protein structure and glycosylation. High-glycoimpact amino acids co-evolve with glycosites, and glycoimpact is high when estimates of amino acid conservation and variant pathogenicity diverge. We report thousands of disease variants near glycosites with high-glycoimpact, including several with known links to aberrant glycosylation (e.g., Oculocutaneous Albinism, Jakob-Creutzfeldt disease, Gerstmann-Straussler-Scheinker, and Gaucher’s Disease). Finally, glycoimpact quantification is validated by studying oligomannose-complex glycan ratios on HIV ENV, differential sialylation on IgG3 Fc, differential glycosylation on SARS-CoV-2 Spike, and fucose-modulated function of a tuberculosis monoclonal antibody. Finally, to test the causality of protein-glycan associations, we created 5 glycoimpact-designed novel Rituximab variants, 4 of which substantially changed glycoprofiles as predicted. In all, we report that site-specific glycan biosynthesis is influenced by underlying protein structure, enabling glycan structure prediction and genetic sequence-guided glycoengineering.

## Introduction

DNA, RNA, and protein synthesis follow DNA and RNA templates. In contrast, glycosylation is understood as primarily regulated by the cellular environment.^3–6^ While glycosylation varies across species, cell types and cell lines, specific glycosites on proteins reproducibly host different glycans, termed glycosite-specific microheterogeneity.^7^ Microheterogeneity suggests local protein structure may also constrain glycosylation.

Many studies have identified specific examples of how primary protein structure or sequence can influence glycosylation patterns. First, the N-glycosylation sequon^8^ defines where glycans covalently attach, i.e., to asparagines (N) with a downstream (N+2) serine (S) or threonine (T) separated by any amino acid (X) except proline (NX[S/T]). Variation at N+1 impacts glycan complexity^9^ and accessibility impacts glycosite occupancy.^10^ Glycosylation of sequons ending in threonine is ∼40 times more efficient than those with serine.^11^ Studies have identified specific examples wherein primary protein structure or sequence can influence glycosylation patterns. Upstream of the glycosite, a phenylalanine (F) to alanine (A) substitution in human IgG3 can increase bi-galactose structures with a core-fucose.^12^ Additionally, influenza evolves hemagglutinin glycosylation sites to facilitate immune evasion.^13^ Glycoprotein-determined glycan evolution, used by HIV for immune evasion, has been leveraged to engineer better vaccine epitopes.^14^ Tools like GlycoSiteAlign^15^ and mutagenesis studies have offered expansions of the founding sequon structure (NX[S/T]) including the enhanced aromatic sequon—an aromatic residue upstream of the glycosite (N-2) that can influence glycan complexity^16^ with a variable impact given the N+1 variation.^17^ Several studies since 2002 have shown variable success in predicting glycosites from sequons and hidden features using machine learning,^18–21^ though none have predicted beyond glycosite presence or the root monosaccharide.

Beyond a protein primary structure (i.e., protein sequence), secondary structures (e.g., β-sheets and α-helixes) can influence glycosylation^22^ while tertiary structures (concavity, accessibility and hydrophobicity) also impact glycosite occupancy.^10^ Crucial bottlenecks in glycan processing such as, ERManI (MAN1B1) and Golgi Mannosidase IA (MAN1A1), have been co-crystalized with a Man9 glycan to study specific glycoconjugate features favoring or inhibiting this interaction.^23–26^ A study of >150 glycoproteins revealed differential glycosylation as a function of protein structure, wherein low accessibility sites impacted core fucosylation and branching;^27^ similar accessibility constraints predicted oligomannose on the SARS-CoV-2 spike protein.^28^ Furthermore, a structural analysis found FUT8 activity is impacted by glycosite depth.^29^ Site-specific kinetics for glycosylation of PDI (Protein Disulfide Isomerase)^30–32^ and five additional glycoproteins^33^ also showed that protein structure influences occupancy. Recent studies have aimed to graft glycans to proteins^20,34–36^, providing valuable tools for exploring mechanistic and steric associations between glycans and folded proteins. Yet, they do not explore generalizable associations between glycan and protein structure features that can be mapped back to the genome.

Here, we identified associations between glycosylation and local protein structures (1D to 3D). We did this by analyzing trends in site-specific glycosylation^37^ and comparing biosynthetic precursors of observed glycans^38^ with local glycoprotein structure features.^39^ We call associations between protein structures and glycan structures ***“Intra-molecular Relations”*** (IMR); IMR are quantified using metrics associating protein structures and glycans structures (e.g., regression models, correlation, or Fisher Exact test) . Further, we call the expected difference in glycan structure following protein sequence or structure changes, ***“Glycoimpact;”*** the difference in IMR associated with protein sequence or structure defining the change. Glycoimpact is detected and validated here by comparison to evolutionary substitution matrices, variant pathogenicity scores, and glycosite co-evolution. Further, glycosite-proximal pathogenic variants correspond to higher glycoimpact substitutions. Finally, glycoimpact accurately predicts changes in glycan complexity, galactosylation, sialylation, and functional glycosylation. These results show IMR represent generalizable associations between protein structure and glycosylation (**Figure 1**). Consequently, IMR suggest protein structure can constrain glycosylation, providing increased clarity of factors impacting microheterogeneity.

**Figure 1.**
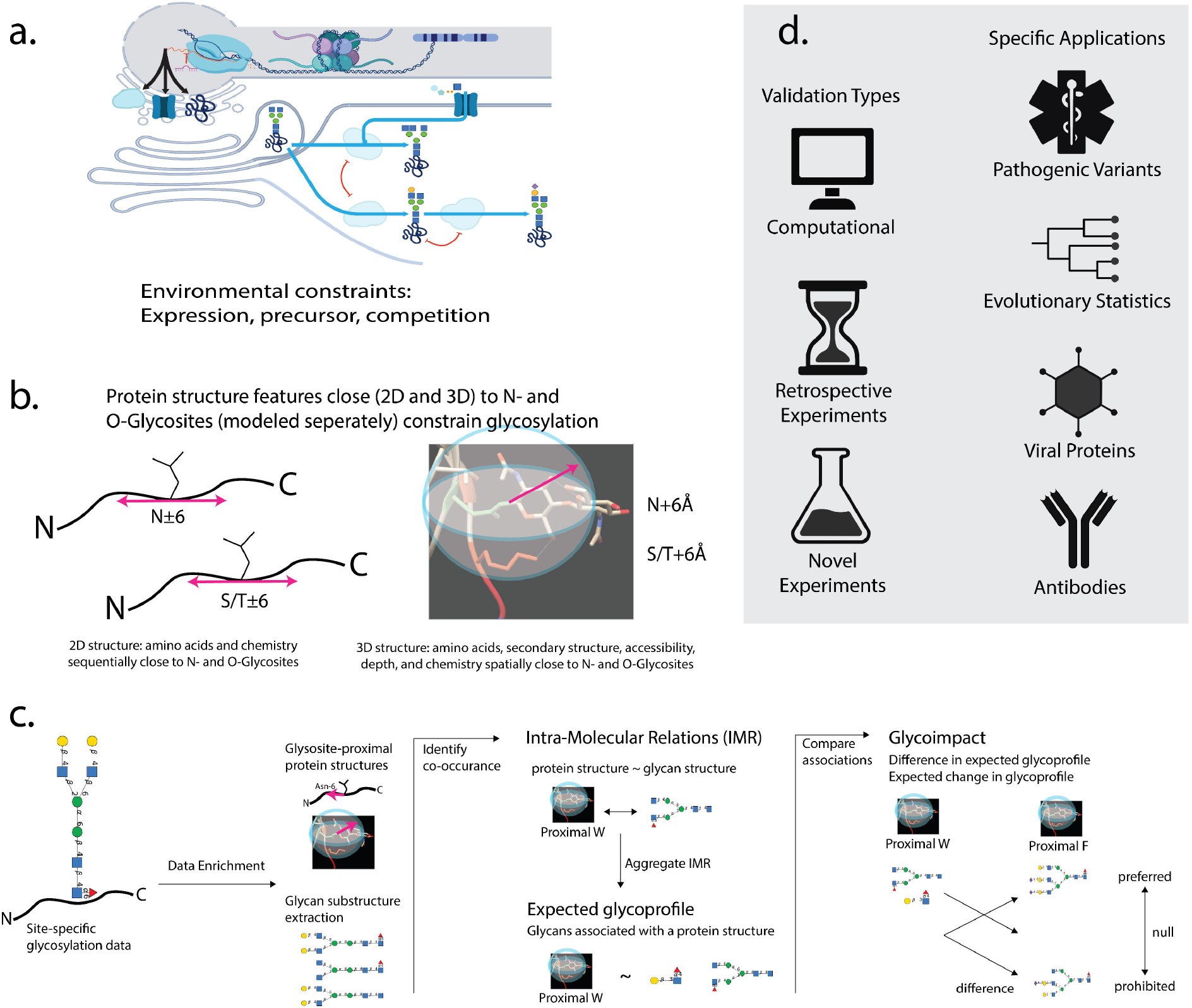
(**a**) DNA, RNA, and protein biosynthesis are template-driven processes. Glycan biosynthesis is described as metabolically and enzymatically constrained. (**b**) Glycoprotein 2D and 3D structural features are considered in relation to proximal glycosylation. (**c**) Site-specific glycosylation data (left) are used to estimate associations between site-specific protein glycan occurrence and proximal sequence (b, left) or structure (b, right), as quantified with our Intra-Molecular Relations (IMR) metric (middle). Following amino acid changes, IMR reflect the agreement between expected glycosylation and a glycoprotein structure, while disagreement is singled out and called “glycoimpact” to capture glycan sequence changes (right). Glycans are represented using IUPAC-extension for glycans and the Symbol Nomenclature for Glycans (SNFG^1,2^); Mannose (Man), Galactose (Gal), Sialic Acid (Neu5Ac), N-Acetylglucosamine (GlcNAc), Fucose (Fuc). (**d**) We validated predictions of glycoprotein structure constraining glycosylation with computational results, published experimental data, and novel experimental results across pathogenic gene variants, evolution, viral proteins, and antibodies.

## Results

### UniCarbKB site-specific glycosylation is representative of human glycosites in PDB

We first tested if glycan structure correlates with protein structure. To do this, we curated a Protein-Glycan Dataset (PGD), a compendium of experimentally-measured and expert-curated site-specific glycosylation on human glycoproteins collected from the UniCarbKB^41^ and GlyConnect^37^ databases (description and URLs, **see Methods**). Glycan structures were decomposed into glycan substructures (i.e., biosynthetic intermediates using GlyCompare^38^). Glycosite-proximal protein structure features were extracted from protein sequence and 3D structure models. “Sequence proximal” was defined as five residues C-or N-terminus to the glycosite and spatial proximity was defined by minimum distance— the minimum 3D Euclidean distance between the nearest atoms in each residue. The resulting dataset includes 111 human glycoproteins (98 glycoproteins with N-glycosylation sites and 38 glycoproteins with O-glycosylation sites) 306 glycosylation sites and 4,263 glycosylation events (3,563 N- and 700 O-linked glycosylation events) (Supplementary **Figure 1**). In this work, we focused training on human glycoproteins and validated predictions to human-expressed glycoproteins. Human and human expressed glycoproteins were used to assert consistency in the underlying biosynthetic machinery.

We verified that annotated glycosites in PGD are representative of all glycosite structures in the human secretome;^42^ all secretome glycosites fall within a dimensionality reduction trained only on PGD glycosite structures (**Figure 2a**, see **Supplementary Results**, Supplementary **Figure 2**). Thus, PGD can support generalizable conclusions about protein-glycan associations.

**Figure 2.**
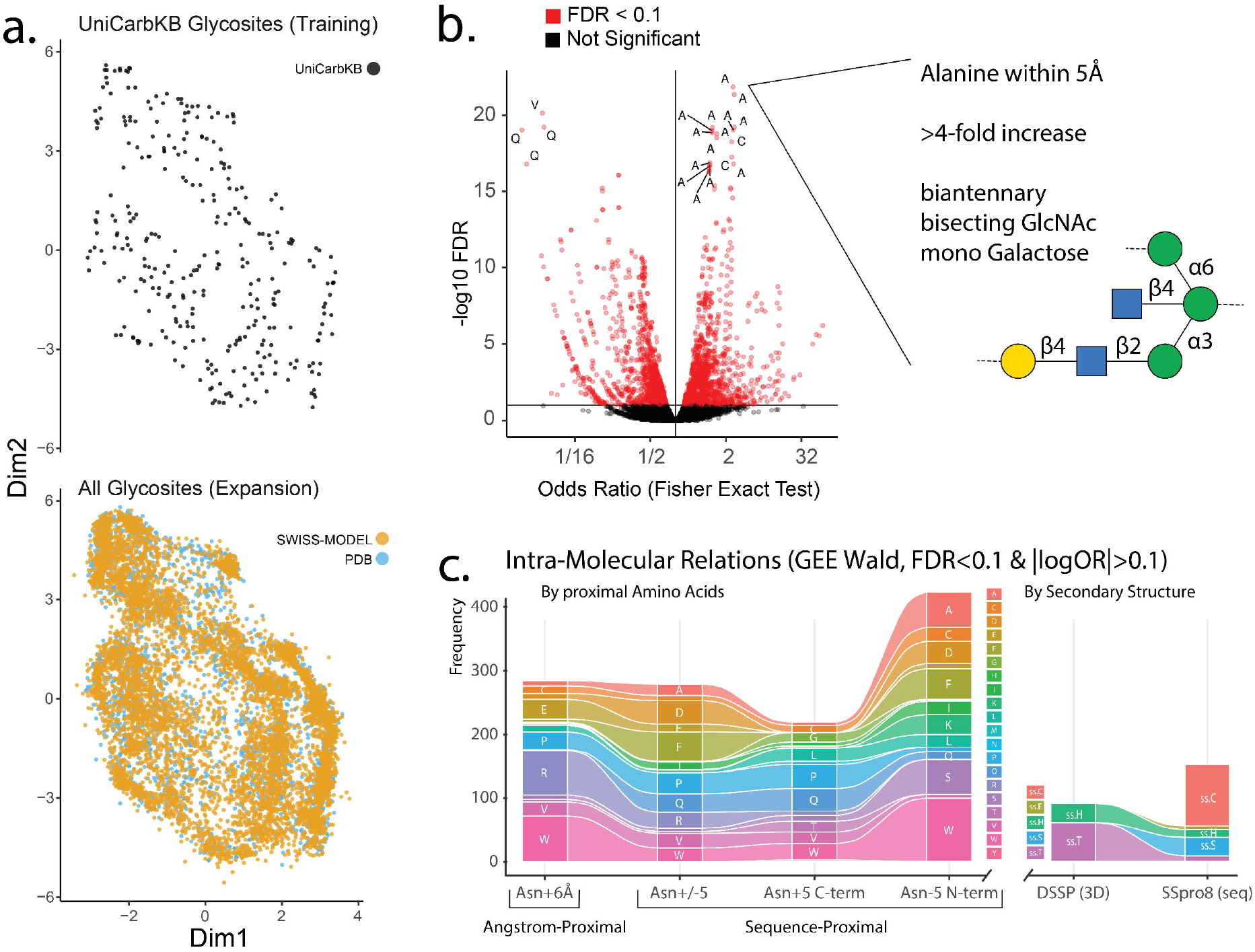
N- and O-glycan substructures associated with glycosite-proximal protein structure. **(a)** Dimensionality reduction trained on UniCarbKB glycosite-proximal protein structures. When projected into the UniCarbKB-trained space, all human UniProt annotated glycosites appear within the UniCarbKB sequence space implying that UniCarbKB glycosites are sufficiently representative of most documented glycosites. The scatterplots show a two-dimensional projection of the FAMD using a Uniform Manifold Approximation and Projection (UMAP).^40^ Each point is a site on a glycoprotein, each color indicates the source of that protein. (**b**) Volcano plot of the log odds ratio and False Discovery Rate adjusted p-values from a Fisher exact test between co-occurring glycan and protein structures. We observe IMR, with most significant relations associated with alanine, cysteine, valine, and glutamine (A, C, V, Q, respectively). (**c**) IMR were also estimated by logistic GEE controlling for protein-bias. The number of significant (FDR<0.1, |log(OR)|>0.1) IMR relating to structurally proximal AAs (N+6Å), sequence proximal AAs C-terminal (N+5), N-terminal (N-5) or either direction (N+/-5), predicted secondary structure from sequence (SSpro8) and structure (DSSP): alpha-helix (ss.H), extended strand (ss.E), beta-bridge (ss.B), turn (ss.T) bend (ss.S), other (ss.C).

### Site-specific glycosylation data contain significant associations between protein sequence, 3D protein structures, and glycosylation

We next examined the associations between glycan substructures (e.g., bisecting GlcNAc) and glycosite-proximal protein features (e.g., proximal tyrosine) within PGD. Substantial and significant (Supplementary **Figure 3a**) associations between glycan substructures and protein features were termed “intramolecular relations” (**IMR**) (Figure 1c). Fisher exact test selected 10,111 IMR with 9,296 positive and 815 negative associations (**Figure 2b**).

### Approximately 20% of IMR predict glycosylation from protein structure with high confidence

The PGD contains several highly deterministic amino acids (AA) specific IMR for which the presence or absence of a glycan substructure is highly determined by the presence or absence of a proximal AA (i.e., Pr(glycan substructure | proximal amino acid) is 0.999-1 or 0-0.001). Overall, confidence in glycan presence increases when a specific protein structure feature is present (Supplementary **Figure 4**). For IMR with a spatially proximal AA (within 6Å), 20.2% are highly deterministic of glycan substructures. Additionally, 32.4% and 17.5% of down- and upstream AA IMR (+/-5 AA) are highly deterministic. The certainty with which an AA predicts a glycan structure decreases substantially when proximal AA are absent (Supplementary **Figure 4a**). The high-confidence IMR count is proportional to the number of unique substructures revealing that these IMR are not dominated by small numbers of glycan motifs, notable substructures (Supplementary **Figure 4b**).

Among the highly deterministic protein-glycan relations (1,725 N-glycosylation events), 1,553 glycans contain a N-acetylglucosamine (GlcNAc) on the α-1,6-mannose branch (Glc2NAc(β1-6)Man(α1-6)Man(β1-4)Glc2NAc), indicative of complex N-glycosylation. All 75 high-confidence IMR with a downstream tryptophan (W) include glycans with the hybrid/complex substructure. These observations suggest that W would be sufficient to escape oligomannose structures. Similarly, in 454 O-glycosylation high-confidence IMR, 237 contain 6⍰-sialyl-N-acetyllactosamine (Neu5Ac(α2-6)Gal(β1-3)GalNAc). Of six events containing a sequence-proximal W, each also contained a sialyl-T antigen. Thus, several IMR link protein features, such as proximal W, with specific glycan features.

### Glycosite-proximal, sequence, structure, and chemistry correlate with glycan structure

To generate testable hypothetical protein structural constraints on glycan biosynthesis, we modeled IMR using hierarchical regression to minimize relevant bias. We quantified specific IMR using univariate logistic generalized estimation equations (GEE) to probe the glycan-protein co-occurrences in PGD and to control for protein identity effects (see **Methods**). A list of GEE odds ratios (OR) (each describing the association between a protein structure and one of several glycan substructures) describes typical glycosylation near a given protein structure; the OR list indicates the “expected substructure abundance/absence” like an “expected glycoprofile” given a proximal protein structure.

We discovered 1,715 significant N-glycan IMR (FDR<0.1, |log(OR)|>0.1), many of which were associated with spatially proximal (N+6Å) and sequence-proximal AA (N+/-5 residues). Stratifying sequence-proximal effects, we found almost twice as many IMR involving upstream (N-5, N-terminal) than downstream (N+5, C-terminal) AA. Among the downstream AA effects, tryptophan (W), alanine (A), serine (S), and F are most impactful (n = 99, 55, 55, and 48 IMR respectively). W also has many IMR when downstream or spatially proximal (n = 26). Spatially proximal arginine (R) (n = 70) and downstream glutamine (Q) (n = 35) are the largest effectors. Finally, glycosylation sites on turns have many IMR (n = 61) (**Figure 2c**).

Turn-associated IMR include >3-fold increases in di- and tri-sialylated tetra-antennary and >2-fold increases in mono- and di-galactosylated structures with core fucose; all positively correlated structures have at least one galactose (Gal) while not all are core fucosylated. Increased complexity at a high-exposure glycosite (i.e., turn) is consistent with prior results demonstrating an inverse association between complexity and depth^27,43^. Structurally proximal Q is associated with a >20-fold increase in monosialylated triantennary structures and a 10-fold decrease in tetraantennary structures (**Figure 3a**). Histidine (H), threonine (T), and valine (V) show correlation with increasing GalNAc[4S] (**Figure 3b**). Expanding on **Figure 3a-b**, we further interrogated IMR for specific protein structure features (i.e., proximal amino acid or local secondary structure) by comparing the IMR odds ratio to monosaccharide count for several monosaccharides (**Figure 3c**). A biclustering highlights at least two major groups of glycan features differentially impacted by different protein structure influences, mirroring the N-glycan/O-glycan dichotomy in glycosylation. Providing clues to the elusive O-glycosylation site, proximal A is negatively correlated with Gal and GlcNAc but positively correlated with N-acetylgalactosamine (GalNAc). Conversely, T and H are positively associated with GlcNAc and Gal but negatively correlated with β-GalNAc. As expected, GlcNAc and complex-glycans follow similar trends to Neu5Ac. However, Neu5Ac, GlcNAc and Gal trends diverge near proline (P), cysteine (C), and V; these amino acids may act as limiters of high-complexity (**Figure 3c**).

**Figure 3.**
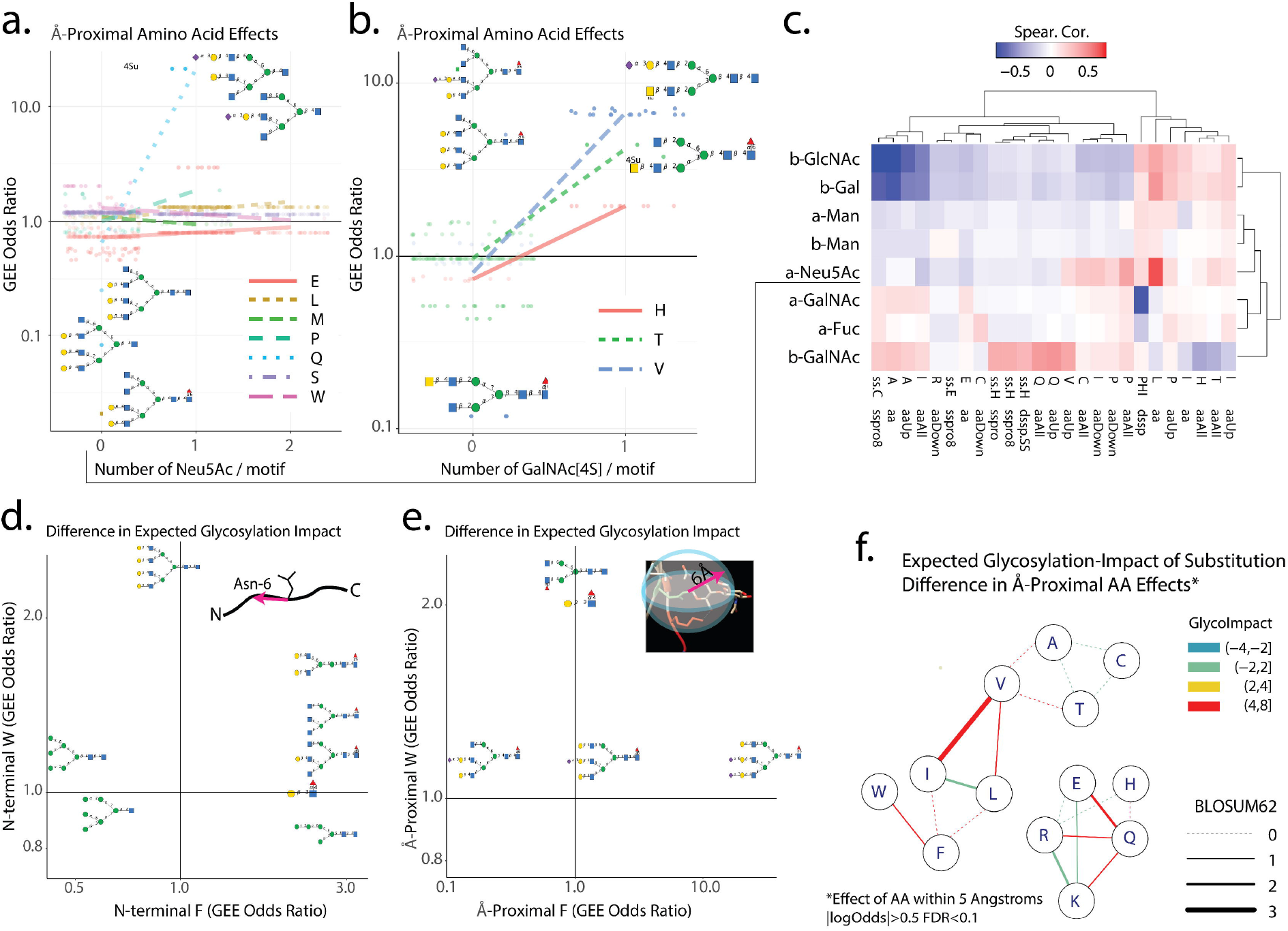
Glycosite-proximal amino acids impact glycosylation. Specific IMRs were discovered by logistic GEE controlling for protein-bias. **(a-b**) IMR (FDR<0.1, |log(OR)|>0.1) relating structurally-proximal amino acids to motifs stratified by the number of Sialic Acids (**a**) and 4-Sulfated GalNAc (**b**). **(c)** Spearman correlation between the monosaccharide count of glycan substructures from protein-structure features; protein structure features with an average absolute correlation > 0.2 were retained. Terms used here are consistent with panel **a** but “aa” denotes a structurally proximal amino acid, “aaUp,” “aaDown,” and “aaAll” denotes N-terminus, C-terminus, or any sequence proximal amino acid. (**d-e**) IMR (FDR<0.1, | log(OR)|>0.1) compared across two sequence-proximal (d) and spatially proximal (**e**) amino acids, phenylalanine (F) and tryptophan (W). The direct comparison of proximal-amino acid effects visualized the expected change in glycosylation associated with that substitution. (**f**) Network depicting the glycoimpact (see **Methods**) of spatially proximal (within 5) substitutions for structurally low impact (BLOSUM62) substitutions. Panel **f** shows glycoimpact predicted from BLAMO 0.5:0.1 (|log(OR)| > 0.5, FDR<0.1), see ***Supplementary* Figure 5** for additional thresholds, raw scores, and sequence-proximal substitution predictions. Note that plots including glycan substructures were manually curated to summarize the selected glycan substructures.

### Amino acid changes can predict glycan structural changes, “Glycoimpact”

The difference in expected glycosylation associated with two distinct amino acids can indicate the expected change in glycosylation following substitution between these amino acids (AA). As mentioned previously, the expected impact of substitution on glycosylation, is termed “glycoimpact.” Such variations are estimated by comparing expected glycoprofiles (see above) for two AA-pairs (e.g. proximal valine and isoleucine). Specifically, glycoimpact is measured as the difference (z-score normalized Euclidean distance) in expected glycoprofiles between each AA-pair observed near glycosites of two protein structures (**Figure 3**, see **Methods**).

Many AA-substitutions are glycoimpactful on glycan biosynthesis, as exemplified by the F to W substitution (**Figure 3d-e, Supplementary Results**). Upstream F is associated with bi- and tri-antennary (>3-fold) while upstream W is associated with tetraantennary terminal Gal (>2-fold) implicating this substitution in branching (**Figure 3d**). Additionally, spatially proximal F is strongly associated with increased sialylation (>10-fold) implicating this amino acid in glycan maturation (**Figure 3e**). These predictions suggest that W to F substitution has a large impact on glycoprofiles. Relevant substitution events are highlighted as “glycoimpactful” (significantly high glycoimpact) and structurally ambivalent (BLOSUM>0) (**Figure 3f**); those high-glycoimpact (red) and low structural impact (thick line) are substitutions of particular note as potential avenues of non-structurally impactful glycan modulation.

### Glycoimpact explains divergence of BLOSUM and PAM substitution matrices

We call the glycoimpact AA-substitution matrix the <u>BL</u>OSUM-<u>PAM O</u>rthology matrix, BLAMO x:y, where x and y refer to the log odds ratio and FDR thresholds respectively. Sub-threshold odds ratios, those insignificant or unsubstantial by the x:y threshold, are excluded from the glycoimpact calculation. **Figure 3f** displays a subset of BLAMO 0.5:0.1 relations. Comparisons can be made at multiple thresholds (Supplementary **Figure** 5) but BLAMO 0.5:0.1 presents as a more representative, normative, and stable threshold.

To further explore the relevance of glycoimpact, we compared it to conservation-based measures of amino acid substitution impact. The PAM^44^ and BLOSUM^45^ AA substitution matrices are popular but distinct estimates. PAM is based on global protein alignments and tends to reflect functional conservation, while BLOSUM relies on local protein alignment reflecting mostly structural conservation (e.g., domains).^46,47^ Thus, PAM and BLOSUM diverge more when protein function is not fully described by protein structure.

Since changes in glycan structure can modify protein function in a protein structure-independent manner, we examined consistency between PAM and BLOSUM estimates across null and high glycoimpact substitutions. Comparing PAM and BLOSUM scores at multiple thresholds (RMSE(PAM_i,j_,BLOSUM_i,j_)), we found that error in 4 of 5 PAM-BLOSUM comparisons was significantly correlated to glycoimpact (BLAMO 0.5:0.1) for high-glycoimpact (GI>2.5) substitutions (**Figure 4a**, Supplementary **Figure** 5). Acknowledging the limited effect size (R<0.4), these associations are strongly significant and consistent across multiple comparisons suggesting, as noted, a significant and global but noisy trend. The correlation between high-glycoimpact substitutions and PAM-BLOSUM inconsistency is maintained for most PAM and BLOSUM thresholds (Supplementary **Figure 6**). Both protein structure and glycosylation are necessary to fully explain protein function, and these results suggest a positive relationship between glycoimpact and the failure of structure (BLOSUM) to completely explain function (PAM). Given this relationship, we refer to the glycoimpact substitution matrix as the BLOSUM-PAM Orthology (BLAMO) matrix.

**Figure 4.**
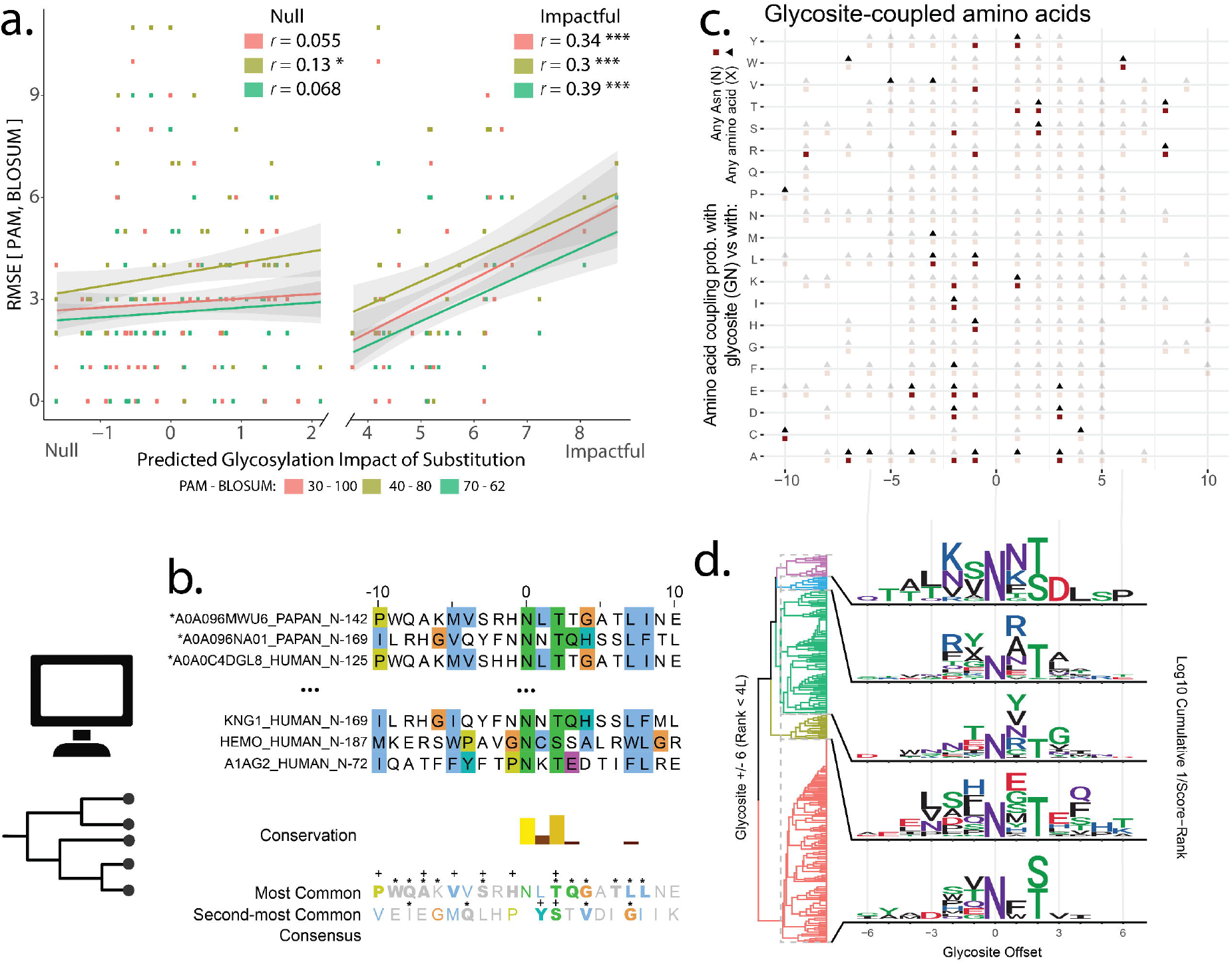
Glycoimpact of amino acid substitution correlates with evolution metrics and conservation. (**a**) Comparison of the error between the PAM and BLOSUM substitution matrices and the glycoimpact for corresponding substitutions. Linear regressions are split into null glycoimpact (<2.5) and glycoimpactful (>2.5). Glycoimpact scores from BLAMO 0.5:0.1 were used; those computed from strong IMR (|log(OR)| > 0.5, FDR<0.1). Error (y-axis) was calculated as the root mean square error between PAM and BLOSUM scores. Pearson’s Correlation (*r*) significance indicated as <0.001 (***), <0.01 (**), and <0.05 (*). (**b**) Glycosite Alignments ^15^ corresponding to GlyConnect-documented tetraantennary structures (Hex:7 HexNAc:6) with no sialic acid or fucose. See ***Supplementary Figure 9*** for full alignment. The first (top line) and second (bottom line) most popular amino acids are displayed for each position N+/-30. Consensus AAs consistent with other analyses are highlighted in bold and marked with a “+” (Glycosite-coupled, **Figure 4**h) or “*” (high-influence AA, **Figure 2**c). (**c**) An aggregation of all residue-glycosite enrichments at N+/-10 (hypergeometric enrichment illustrated in panel ***Supplementary Figure 7****c*). The proportion of high-ranking Evolutionary Couplings (EC) for each amino acid (rows) at the column-specified relative position was compared with glycosites (GN), asparagines (N), or any residue (X). An opaque red circle indicates that for the residue (row) in the given position (column), high-EC proportion is higher with GN than N. An opaque black triangle indicates high-EC proportion is higher with GN than X. A transparent circle or square indicates GN was not significantly more coupled (hypergeometric test). Significance was assessed at multiple EC-rank thresholds between L/3 and 3L (see Methods) and pooled using Fisher’s method (FDR <0.1). (**d**) Hierarchical clustering of coupling-masked (Rank<4L) amino acids surrounding a glycosite (+/-6 AA). Each of 5 clusters was summarized as a motif. Height is the log of cumulative reciprocal EC-Rank with a pseudo-count of 0.25. The asparagine at the center was fixed at 2 for context. Residues are colored by chemical properties. A more granular clustering, 25 clusters is included in the supplement (***Supplementary Figure 8***).

### High glycoimpact residues are conserved around N-glycosites

If glycosite-proximal amino acids influence functional glycosylation, there should be evolutionary pressure imposed beyond the classical NX[S/T] N-glycan sequon. To map the broader glycosite structure, we aligned the surrounding sequences (five AA upstream and downstream) of N-glycosites (Figure 4b-c) and examined conservation and evolutionary coupling (EC). EC-derived from 2,005 glycoprotein alignments (see **Methods, Supplementary Results**),^48^ substantiates enrichment between N-glycosites and flanking residues (Supplementary Figure 7a). We also observed several position-specific glycosite-coupled residues including S and T at N+2, F at N-2, and tyrosine (Y) at N-1 (pooled hypergeometric, FDR<0.1, Supplementary **Figure 7**c, **Figure 4c**). These findings are consistent with previous observations of the original sequon and enhanced aromatic sequon.^17^ Glycosite-coupled residues are also consistent with IMR-observed high-impact residues (**Figure 2c**). Of the ten high-impact upstream residues, seven show enriched glycosite-coupling when they appear upstream. To further highlight global glycosite structure, we clustered glycosites using coupling probabilities (**Figure 4d**, Supplementary **Figure** 8, see Methods). The N+2 aspartic acid (Asp) enriched in the univariate analysis (**Figure 4c**) co-occurs with an N-2 lysine (K) (Figure 4d, motif 1). Alternatively, glutamic acid (E) is more likely to co-occur with other E residues (N -4, +1, and +3) with an N+2 T-containing sequon (**Figure 4d**, motif 4). These couplings are reflective of evolutionary pressure surrounding the glycosylation sites.

We next aligned^15^ glycosites permitting a tetraantennary N-glycan lacking fucose or sialic acids (**Figure 4b**). We examined the glycosite alignment for consistency with high-influence AA (Figure 2c) and glycosite-coupled residues (**Figure 4c**). Of 20 glycosite-flanking AA, 16 show consistency between the first or second most common AA and either the high-influence or glycosite coupled residues (see **Supplementary Results**). At nearly every glycosite-flanking residue (N+/-10) there is consistency between these three analyses, further corroborating that protein structure constrains glycosylation.

### Glycoimpact correlates with discrepancy between functional variant predictions

Dozens of algorithms predict functional and pathogenic effects of genetic variants,^49–51^ incorporating information ranging from sequences to protein structure, thus sometimes reporting different variants as deleterious. We hypothesized the differences in some pathogenicity scores between algorithms could be explained by glycoimpact (BLAMO 0.5:0.1, see **Methods**). Across 3,549,910 nonsynonymous mutations, we measured the disagreement (RMSE) between each of 27 rank-normalized functional impact prediction tools, precomputed with dbNSFP;^49^ pathogenicity score divergence was then correlated with glycoimpact. After hierarchical clustering on the divergence-glycoimpact correlation coefficients, pathogenicity estimates separated into two major clusters: one containing nearly all (6/7) tools leveraging protein-structure and the other primarily containing conservation, sequence, and/or epigenetic-based tools (**Figure 5b**). Nearly all variant pathogenicity score differences across the two clusters correlated with glycoimpact. These correlations and clustering structure disappear when glycoimpact scores are shuffled (Supplementary **Figure 10**); thus, like BLOSUM-PAM discrepancies, glycoimpact correlates with discrepancies between conservation-based and protein structure-based pathogenicity estimates. These observations further implicate glycosylation as a potent functional regulator and glycoimpact as an appropriate proxy for the importance of glycosylation.

**Figure 5.**
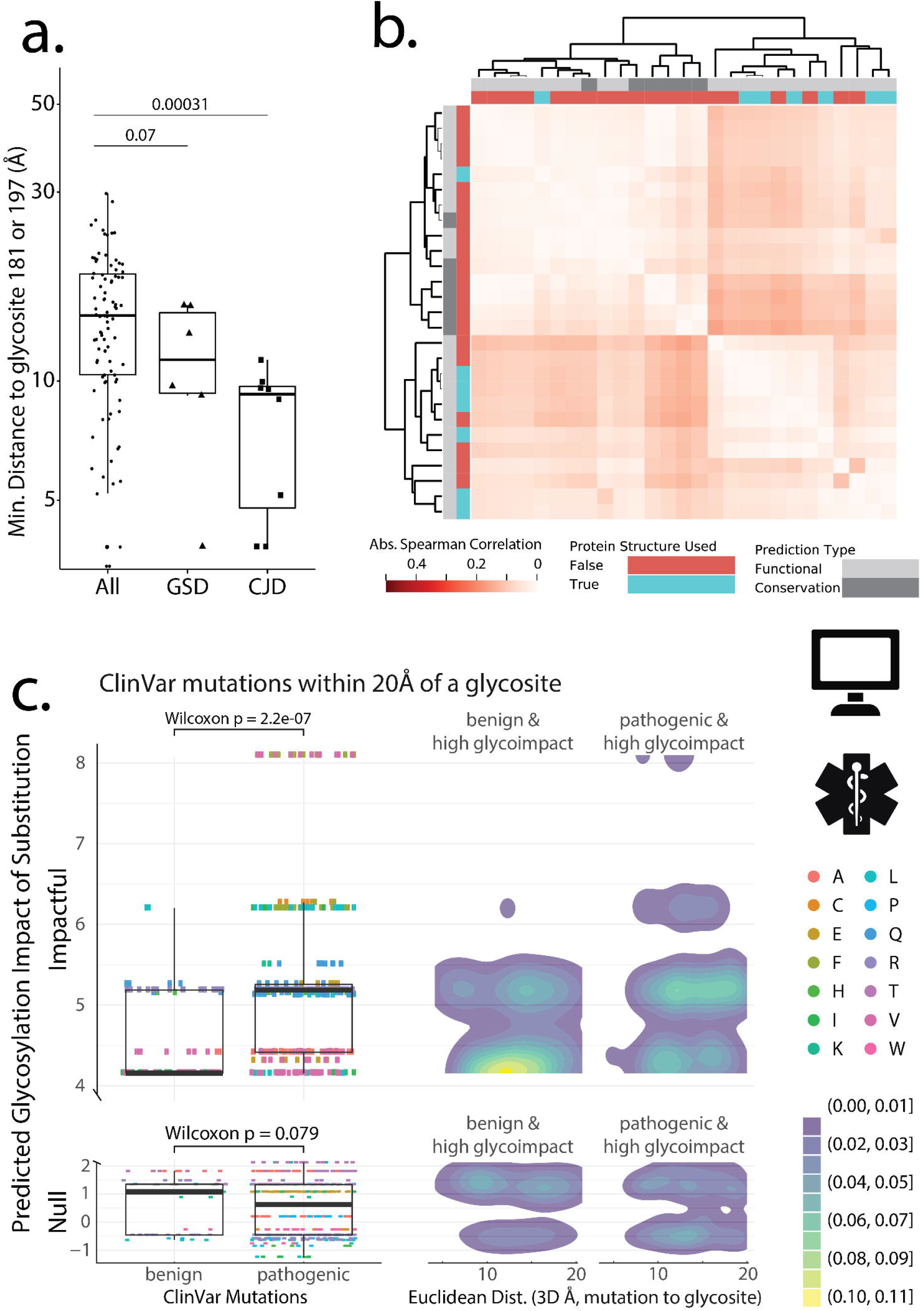
Substitution glycoimpact predicts pathogenicity. (**a**) Boxplots show the min-distance from all residues within human PrP to the N197 or N181 glycosylation sites. Residues are stratified by all sites (All) and causative mutations of prion disease including Creutzfeldt-Jakob disease (CJD) and Gerstmann-Straussler disease (GSS), (one-sided Wilcoxon test). For glycosite-specific proximity, see ***Supplementary Figure 11***. (**b**) A hierarchical clustered heatmap (average-linkage with Euclidean distance) of Spearman correlation coefficients between glycoimpact (BLAMO 0.5:0.1) and error between variant impact prediction scores. Prediction-type and protein structure indicate the training data used to build various tools as described in dbNSFP ^49^. Each row and column refer to a variant function prediction tool. (**c**) Null and impactful glycoimpact (BLAMO 0.5:0.1) stratified by variant pathogenicity in ClinVar within 20Å (min-distance) of an N-glycosylation site. Two-dimensional density plots compare glycoimpact and the glycosite-mutation distance.

### Glycoimpact proposes mechanisms for pathogenic variants in ClinVar and PrP

Because glycoimpact correlates with evolutionary, structural, and functional AA-substitution metrics, we hypothesized it is also pathologically relevant. Thus, we compared glycoimpact and clinical impact for ClinVar-annotated variants within 15Å, 20Å, and 30Å of UniProtKB-annotated glycosites. For all three distances tested, high glycoimpact (BLAMO 0.5:0.1) variants in ClinVar were robustly and significantly higher (Wilcoxon p=2.6e-4, 2.2e-7, and 1.6e-10) for pathogenic variants close to glycosylation sites compared to glycosite-proximal benign variants (**Figure 5c**). Similarly, glycoimpact is higher for likely-pathogenic variants and variants of unknown significance near glycosites.

Examining specific ClinVar-annotated variants, we identified multiple variants in glycosylation-related diseases. For example, Tyrosinase:A355V (P14679) is a high-glycoimpact and glycosite-proximal causal variant in oculocutaneous albinism.^52,53^ While tyrosinase:A355V has not been examined for aberrant glycosylation, deglycosylation disrupts tyrosinase function consistent with type 1 oculocutaneous albinism.^53–57^ Therefore, tyrosinase:A355V may act through aberrant glycosylation. We also observe high-glycoimpact, glycosite-proximal (<30Å), pathogenic (ClinVar) variants in multiple other glycan-modulated diseases including prion diseases,^58,59^ lysosomal storage disorders,^60,61^ and Gaucher’s disease.^62–65^

More broadly, of 1,228 non-benign ClinVar annotated variants on glycoproteins, 340 are high-glycoimpact (GI>2.5) and closer than 30Å to a glycosylation site (**Table 1**, Supplementary **Table 1**, **Supplementary Dataset 1**). This includes major diseases not typically considered related to N-glycosylation including cystic fibrosis, long QT syndrome, renal cell carcinoma, acquired immunodeficiency syndrome, and multiple blood coagulation factor deficiencies. Notably, approximately 36% of the non-benign ClinVar-annotated glycoprotein variants we examined may be impacted by aberrant glycosylation, a potentially underappreciated mechanism of pathogenesis.

**Table 1.**
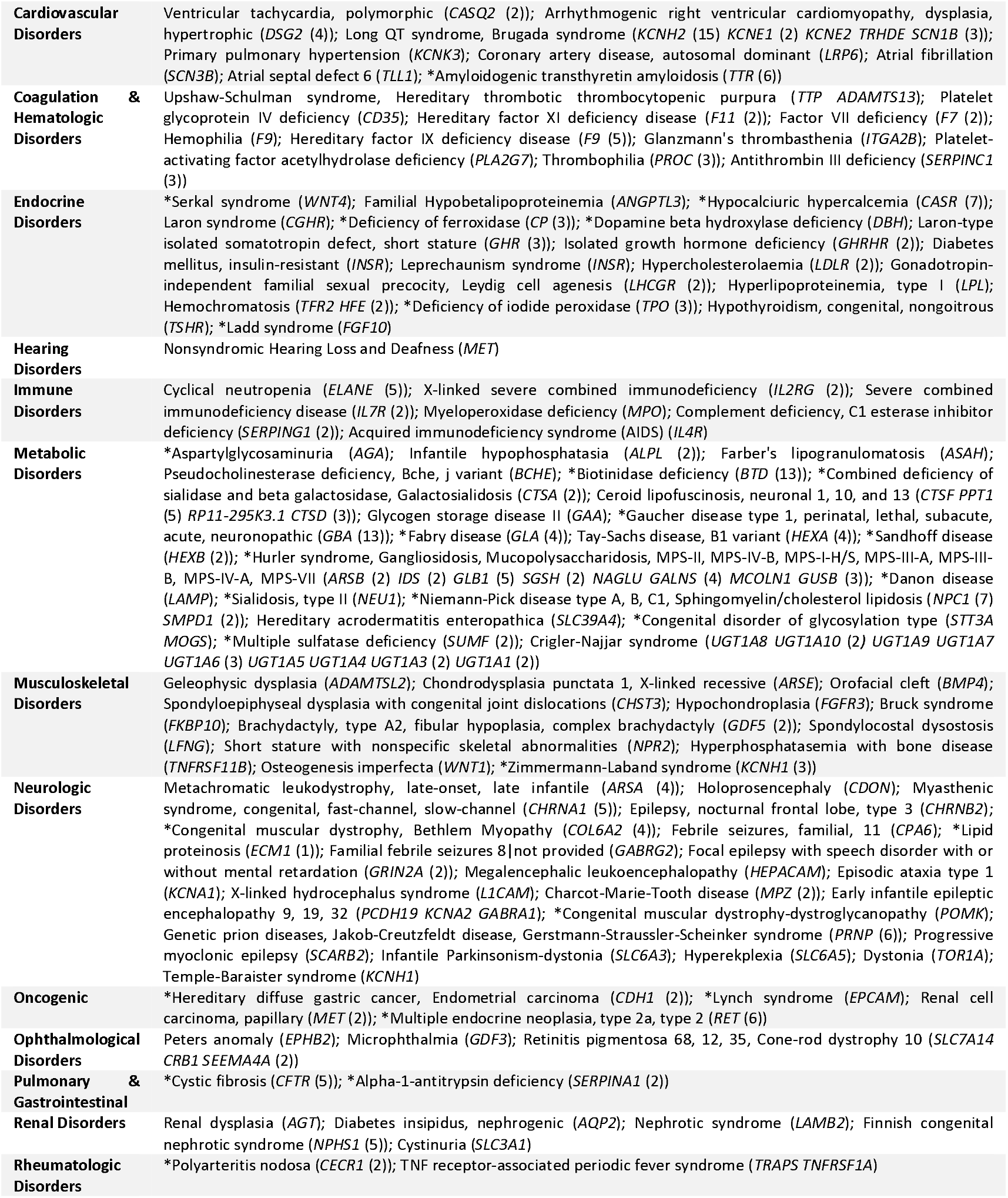
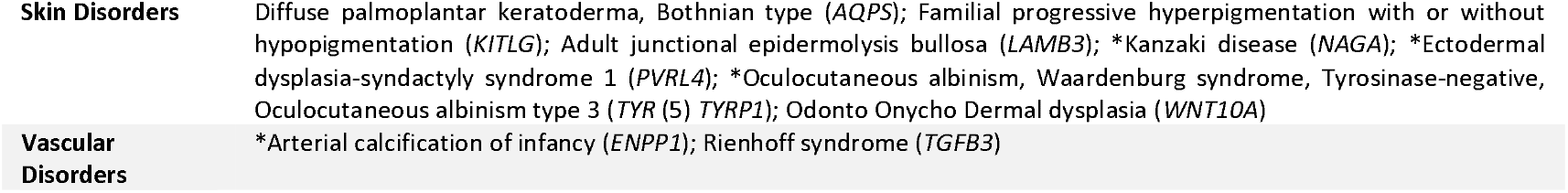
Summary of variants by disease type where variants are high glycoimpact, close to glycosylation sites, and annotated in ClinVar as non-benign. Each specific disease is listed followed by gene name(s) and an integer indicating the number (if larger than 1) of variants in that gene corresponding to each specific disease. “*” indicates a multisystemic disorder.

We further analyzed glycosite proximity among causal variants in prion disease. We measured 3D Euclidean min-distance from the two PrP glycosylation sites, N181 and N197, to all residues in human prion protein (PrP) (including variants causing Creutzfeldt-Jakob disease (CJD)^66,67^ and Gerstmann-Sträussler-Scheinker disease (GSS))^68,69^ (Figure 5a, Supplementary **Table 2**). CJD-causing variants were approximately twice as close to glycosylation sites than the background distribution of all PrP sites (One-sided Wilcoxon p=0.0003). GSS-causative variants also trend closer to glycosites (One-sided Wilcoxon p=0.07) (Figure 5a, Supplementary **Figure 11a-b**). Low expression mutants, an indication of possible aberrant glycosylation, trend closer to site N180 (One-sided Wilcoxon p=0.16) and appeared further from N197 (One-sided Wilcoxon p=0.04) (Supplementary **Figure 11c-d**).

### Protein structure correlates with glycan complexity in HIV gp160

To further validate our predictions, we compared IMR to glycosylation on specific glycoproteins. We first found consistency between previously measured IMR (**Figure 2b**) and site-specific glycan complexity measurements in HIV ENV gp160 (**Figure 6b**).^70^ PGD-measured GEE-estimated IMR suggest that downstream Q was significantly (FDR<1e-8; OR<0.5) predictive of complexity while spatially proximal P and K were weaker but significant distinguishers (FDR<1e-3, FDR<0.1 respectively, **Figure 6a**). As predicted, gp160 glycosites with proximal P (min distance < 6Å) present more oligomannose (Two-sided Wilcoxon p=0.0033), whereas C-terminus-proximal Q glycosites have more complex glycans (Two-sided Wilcoxon p=1e-4, **Figure 6c**). Spatially proximal K, while less significant (**Figure 6c**), had a nonlinear impact on glycan complexity in HIV gp160; first increasing with one proximal K then decreasing with two. The two most significant IMR predicted from PGD (spatially proximal P and C-terminal Q) were consistent with the site-specific glycosylation observed in HIV gp160.

**Figure 6.**
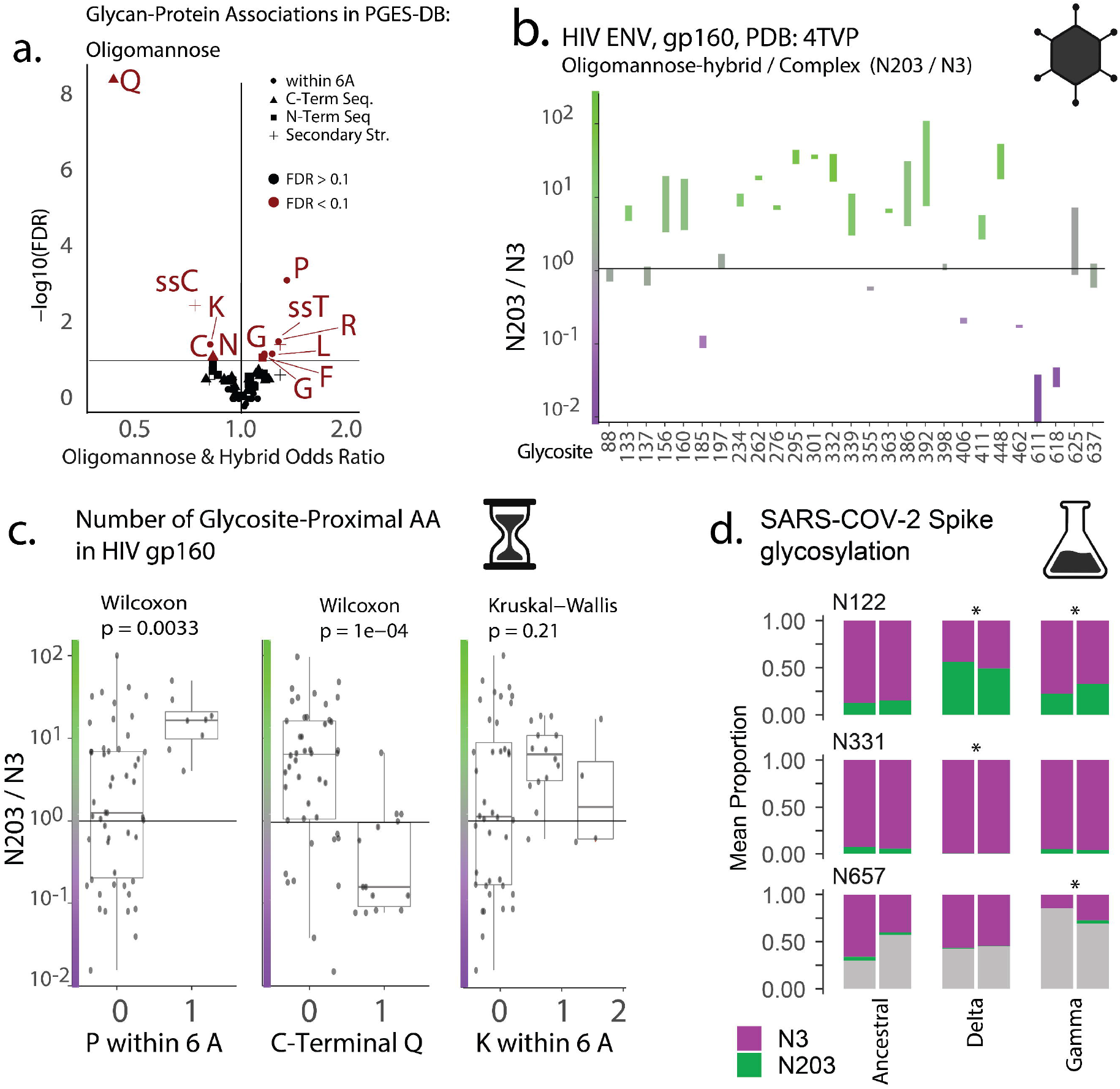
Changes in high-mannose, hybrid, and complex glycosylation are predicted by glycosite-proximal high-glycoimpact substitutions in HIV, SARS-CoV-2. (**a**) GEE-calculated IMR from PGD denoting relations between sequence (triangle & square) or structural protein features (circle & plus) and motifs containing >3 mannose (hybrid & high-mannose). (**b**) The range of mass spectrometry peaks consistent with either oligomannose or hybrid glycans (N203) or complex glycan peaks (N3) at each site on the HIV envelope gp160 (BG505 SOSIP.664, PDB:4TVP).^70^ Relative abundance is represented as a ratio of oligomannose-hybrid to complex glycan peaks (N203/N3). (**c**) Distributions of oligomannose-hybrid to complexity (N203/N3, panel **b**) glycan peaks stratified by proximal protein structure features selected in panel **a.** For the N203/N3 distribution of select IMR, see ***Supplementary Figure 12***. (**d**) Mean proportion of mass-spectrometry-observed peptide with mass offsets corresponding to complex glycan peaks (+3, purple), oligomannose or hybrid glycan peaks (+203, green), or no glycosylation (+0, grey) in the SARS-CoV-2 S1 subunit at 3 sites with significant glycosylation differences between the original ancestral strain, and the Delta and Gamma variants (see ***Supplementary Figure 13*** for all sites). Peak count for each glycosylation offset type (+3, +203, or +0) for each glycosylation site is divided by the total number of peaks for that site to determine the site-specific proportion of each glycosylation type. Significant differential glycosylation (FDR<0.05) in VOC compared to ancestral is indicated by “*”.

### IMR predict differential glycosylation on the SARS-CoV-2 spike glycoprotein

We also predicted glycosylation on SARS-CoV-2 spike protein S1 subunit in the ancestral strain to the Gamma and Delta variants. Several of the >20 glycosylation sites^71,72^ have been implicated with stability, target engagement, furin cleavage, and immune evasion.^72–76^ We found multiple glycosite-proximal mutations within 15Å (min-distance, cubic (3D) expansion of the 6Å IMR training threshold). Gamma spike S1 contains multiple mutations close to glycosylation sites including N17 (L18F, T20N, & D138Y), N61 (P26S & R190S), N122 (L18F, D138Y, & R190S), N616 (D614G), and N657 (H655Y). In the Delta S1, N17 and N122 have 5 and 4 proximal mutations (relative to ancestral S1), respectively, while N165 and N616 each have one high-proximity mutation. Of the glycosite proximal substitutions, only L18F in Gamma and F157V in Delta have a high predicted glycoimpact by IMR. L18F appears within 15Å of N17, N74, N122 in Gamma. Similarly, F157V appears within 15Å of N17, N122, and N165 in Delta. High-impact substitutions appear close to N17 and N122 in both variants.

We measured HEK293-expressed SARS-CoV-2 S1 variant glycan heterogeneity using DeGlyPHER, a mass spectrometry (MS)-based glycoproteomics method,^77^ to determine glycan state and occupancy at each glycosite. We performed two independent technical replicate analyses of the S1 subunit comparing Ancestral to the Gamma and Delta variants and examined 11 of the 12 canonical S1 glycosites (N17 was excluded due to variable signal peptide trimming). Site-specific unoccupied, complex, and oligomannose/hybrid proportions were compared between variants using a Mann-Whitney U test.^77^ P-values were pooled across the two independent replicate analyses using the Fisher method (FDR-corrected). We observed three significant differential glycosylation events (**Figure 6d**). Complex glycans observed at N122 in Ancestral S1 were converted to more oligomannose/hybrid in both Delta (oligomannose/hybrid observations increased nearly 4-fold from 13.9% in S1 to 52.5%; FDR=3.3e-9) and Gamma (oligomannose/hybrid observations nearly doubled to 27.6%; FDR=7.9e-4) variants. Complex glycans at N331 increased marginally from Ancestral S1 in Delta variant (from 93.7% to 99.7%; FDR=0.031). Complex glycans seen at N657 in Ancestral S1 decreased in Gamma variant (by over 2-fold from 53.3% to 21.1%; FDR=1.27e-3). The Gamma S1 monomer was consistently expressed with two novel complex glycosites at N20 and N188.

Based on proximal high-glycoimpact substitutions, we predicted changes at N17, N74 (Gamma only), N122, and N165 (Delta only). Three of four (N122 in Gamma and Delta, N657 in Gamma) predicted differential glycosylation events were consistent with the four observed changes (sensitivity=0.75). Meanwhile, 15 sites where no change was predicted, were consistent with the 17 sites where no change was observed (specificity=0.88). This correct prediction of differential glycosylation is most substantial at N122 in the S1 monomer providing a proof-of-concept that motivates further inspection of differential glycosylation on the more physiological spike trimer and whole virus. This can provide further insights into how differential glycosylation may participate in immune evasion by SARS-CoV-2 and other viruses.

### Glycosite-proximal variation in IgG sequence predicts differential Fc N-glycosylation

We next predicted differential glycosylation for the *ighg1* missense mutation (F299I) in the IgG1 heavy chain.^78^ IgG1 glycosylation impacts both adaptive humoral response^79–82^ and monoclonal antibody (mAb) response.^83–86^ Experimentally, the IgG1:F299I substitution shows a strong glycoimpact. The IgG1 variant expressed in C57BL/6 and BALB/c mouse strains shows less sialylation and digalactosylation compared to similarly expressed *wt* IgG1.^87–89^ Additionally, BALB/c-expressed IgG1:F299I glycosylation is more similar to C57BL/6-expressed IgG1:F299I glycosylation than to glycans on *wt* IgG1 expressed in the same BALB/c animals (**Figure 7b-c**). Fc N-glycans on IgG1:F299I expressed in both BALB/c and C57BL/6 animals have more agalactosylation (Mann-Whitney p=1.02e-6), less digalactosylation, and less mono-, di- and total sialylation (Mann-Whitney p<0.0073) compared to *wt* IgG1 expressed in the same animals (Figure 7c, Supplementary **Table 3**). The increase in galactosylation in *wt* IgG1:F299 is consistent with PGD predicted IMR for upstream (N-terminal) F (**Figure 7a**). Upstream F is associated with increased di-galactosylated biantennary structures (OR>2), while upstream isoleucine (I) is associated with tetraantennary galactosylation. Since only biantennary structures are generally permitted on IgG, the galactose-promoting function of upstream isoleucine should be unrealized in this glycoprotein. The increased sialylation in *wt* IgG1:F299 is also consistent with PGD IMR which show an association between structurally proximal F and disialylated structures (OR>10). These results suggest that glycoimpact can accurately predict several specific glycan epitopes.

**Figure 7.**
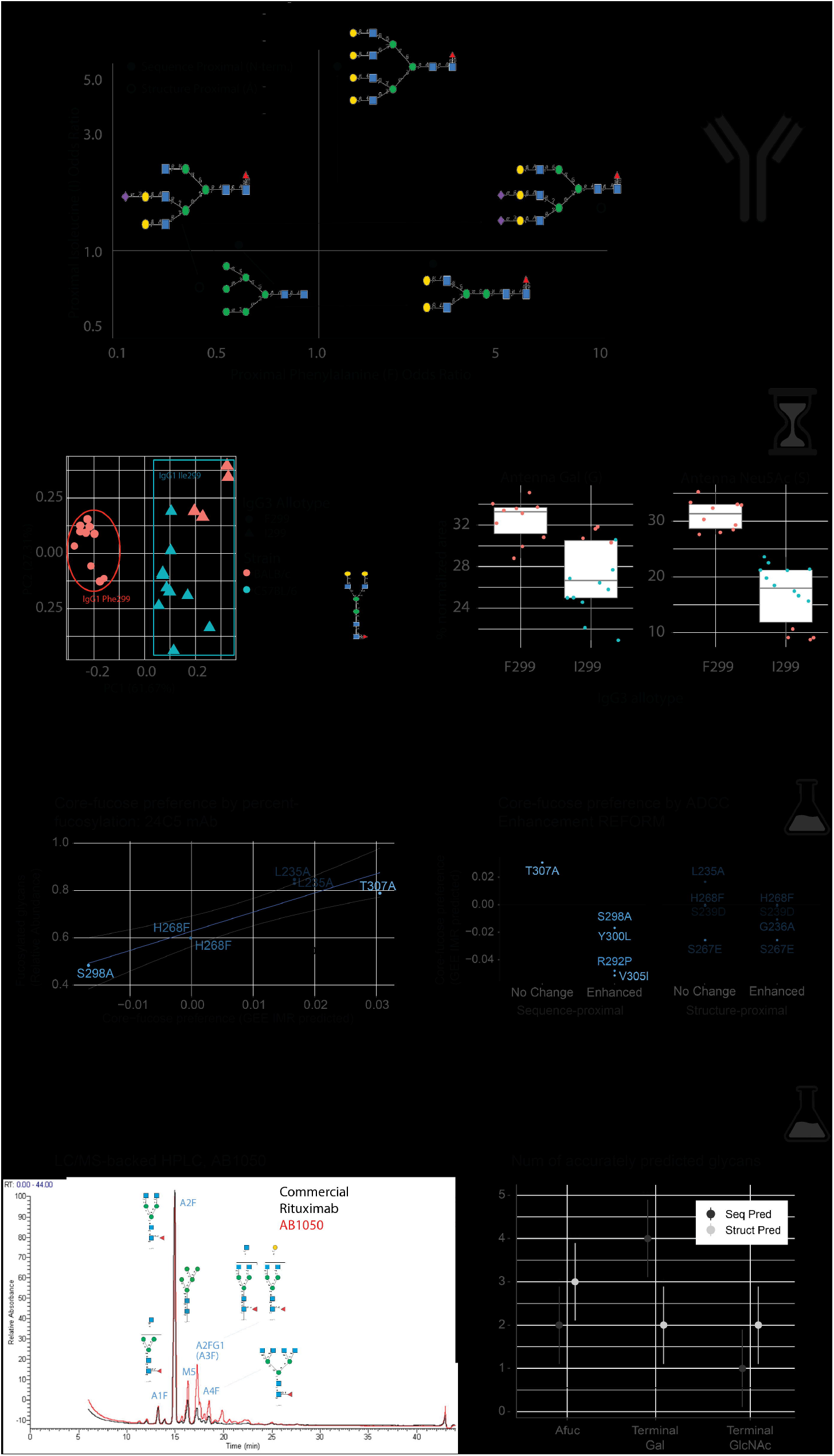
Differential glycosylation is predicted by high-glycoimpact events near glycosylation sites in IgG (**a**) GEE-learned IMR relating to sequence-proximal (upstream/N-terminal) effects of I and F. IgG allotypes, F299 and I299, segregated by Principal Component Analysis of relative abundance (**b**) separate by allotype, not strain. (**c**) Galactose (Gal), Sialylation (Neu5Ac) and Bisecting GlcNAc abundance distributions for IgG1 F299 and I299 allotypes across BALB/c and C57BL/6 mice (see ***Supplementary Figure 14***). (**d-e**) Core-fucose preference (see **Methods, *Supplementary Figure 15***) for each glycosite-proximal variant in the REFORM Fc variant panel.^90^ Variants are colored by index. (**d**) Variants are stratified continuously (top) by percent of fucosylated Fc-cleaved glycans observed on the 24C5 monoclonal antibody. (**e**) Variants are stratified categorically (bottom) by association with enhanced ADCC.^90^ Variants appearing in Fc regions associated with both no-enhancement and enhanced ADCC are shown as transparent. Only L235A and G236A are uniquely associated with ADCC. (**f**) UPLC chromatogram of PNGaseF-released mAb Fc glycans. The expected Rituximab glycoprofile (black) and GlycoTemplated Rituximab variant AB1050 (red) are shown to visualize the changes to the glycoprofile (glycans verified via LC-MS). (**g**) Number (n) of Rituximab variants (of 5) correctly predicted (n) to increase, decrease, or not-change fucosylation, terminal galactose (gal) or terminal N-acetylglucosamine (GlcNAc). Predictions were either from Random estimation, sequence-based IMR-guided predictions (Seq Pred), or protein structure-based IMR-guided predictions (Struct Pred). Significance of consistency between experimental observations and predictions was calculated by a binomial distribution (* < 0.05, ** < 0.01, *** < 0.001)

### Core-fucose preference is associated with percent fucosylation and high-ADCC

To further demonstrate IMR accuracy and functional importance, we compared predicted core-fucose preference to observed fucosylation abundance and ADCC in a Fab-constant Fc-variant antibody panel^90^ (**Figure 7d-e**). Core-fucose preference was calculated as the preference of variant compared to the wild type for N-glycan core motif with or without a core fucose, i.e., Man(b1-4)GlcNAc(b1-4)[Fuc(a1-6)]GlcNAc(b1-4)-Asn and Man(b1-4)GlcNAc(b1-4)GlcNAc(b1-4)-Asn, respectively (see **Methods**).

We compared core-fucose preference for Fc-variant glycosylation on a surface-binding *Mycobacterium tuberculosis*-specific monoclonal antibody (clone 24c5) to demonstrate the accuracy of IMR-based predictions. We determined core-fucose preference predicted from GEE-derived IMR (see **Methods**) for multiple Fc-variants. We then compared predicted core-fucose preference to the capillary electrophoresis measured relative abundance of fucosylated glycans. The Fc-variants naturally stratified into fucose-saturated (fucosylated glycans > 90%) and unsaturated variants (Supplementary **Figure 16**). Fucose saturated variants showed no significant association with predicted fucose preference. However, fucose-unsaturated variants alone (**Figure 7d**) show a strong correlation between predicted core-fucose preference and percent fucosylation (R=0.9, p=0.013). 24c5 Fc glycosylation profiles therefore demonstrate another instance of glycan predictability.

To determine if predicted changes in glycosylation can also predict glycan-modulated behaviors, we examined our ability to predict increases in antibody-dependent cellular cytotoxicity (ADCC) from predicted decreases in core-fucosylation, a well-characterized property of therapeutic antibodies.^86^ We identified 10 glycosite-proximal variants (**Figure 7e**) from the REFORM Fc variant panel^90^ and stratified the ADCC-enhancing variants from those associated with no change in ADCC. Of the five sequence-proximal variants, the non-enhancing variant (T307A) shows a positive preference for core-fucose while all four ADCC associated variants have a negative core-fucose preference. Among spatially proximal variants, several substitutions occur in multiple variants and are not uniquely associated with ADCC enhancement or non-enhancement (greyed out, **Figure 7e**). Of the two-remaining spatially proximal variants, the non-enhancing variant (L235A) shows a positive preference for core-fucose while the ADCC-associated variant (G236A) has a negative core-fucose preference. These ADCC associations with core-fucose preference are consistent with percent fucosylation in 24c5 Fc. T307A and L235A are highly fucosylated in 24c5, predicted to prefer core-fucose, and do not enhance ADCC. Conversely, S298A is least fucosylated in 24c5, shows a negative preference for core-fucose, and is associated with ADCC enhancement. Our predictions therefore recapitulate the immunological implications of allotype-driven differential glycosylation.

### IMR-guided novel Rituximab variants are predictably and differentially glycosylated

To demonstrate our genetic control over glycosylation, we designed 5 Rituximab variants. Fc-variants were created using IMR-designed single amino acid substitutions to modulate branching and fucosylation. Rituximab variants were transiently expressed in 1mL cultures of ExpiCHO cells, purified using affinity chromatography and glycoprofiled using LC-MC-backed UPLC. By creating novel Fc variants we can explore a maximally *ab initio* design space to avoid confounding influence from unrelated design objectives used to create pre-existing Fc variants.

We observed a profound change in glycoprofiles for 4 of the 5 Fc variants including a substantial increase in branching in AB1050 (**Figure 7f**); increased branching is a notable achievement as Fc glycans are almost exclusively biantennary. To better understand the robustness of IMR predictions we compared our structure- and sequence-based predictions to random predictions measured against a binomial distribution. Both structure- and sequence-based predictions performed significantly better than random at predicting fucosylation, and terminal GlcNAc or Gal (**Figure 7g**). Both sequence- and structure-based fucosylation predictions performed significantly better than random (structure- and sequence-based performance were not significantly different from each other). In all, IMR can design completely novel variants to substantially transform glycoprofiles by modifying single residues.

## Discussion

Here, we identified constraints on glycan biosynthesis through the analysis of site-specific protein features^39^ and glycan substructures^38^ and then used these learned associations to effectively engineer glycosylation. With our Protein-Glycan Dataset (PGD), we enumerated and quantified intramolecular relations (IMR) between protein and glycan structure in human glycoproteins. We then computed the expected differential glycosylation, the “glycoimpact,” associated with specific protein structure changes. We validated the importance of glycoimpact by comparison to substitution matrices, evolutionary couplings, and pathogenicity scores. We further examined glycoimpact sensitivity and specificity by comparing predictions with observed glycosylation on PrP, HIV gp160, and IgG glycoproteins. The existence and utility of glycoimpact suggests that glycosylation is constrained by protein structure. We anticipate these constraints depend on which glycosyltransferases can dock and continue glycosylation resulting in a range of feasible or preferred glycosylation for specific glycosites.

The relationship presented here between glycan and protein structures seen in PGD suggests that protein structure provides important information regarding glycan structure. Of course, in extreme expression systems, protein-based predictions of glycosylation and microheterogeneity breakdown as proteostatic stress increases, the unfolded protein response is triggered, and glycosylation machinery fails to meet opportunity for glycosylation.^91,92^ Such low fulfillment is seen in high-yield expression systems^93^ necessitating, in some cases, supplementation with maturing glycoenzymes.^94–96^ Low fulfillment is not inconsistent with protein-based constraints, rather it is a condition which subverts those constraint.

Our quantified protein-glycan IMR describe an expanded glycosite structure, beyond the original sequon definition (NX[S/T]).^8^ We discovered many new sequence-proximal AA IMR both upstream and downstream of traditional glycosites. The enhanced aromatic sequon (EAS)^17^ corroborates one such sequon-expanding IMR. Upstream phenylalanine IMR predicts an increase in Man7 structures (i.e., N-glycan with seven-mannoses) and a decrease in Man6 structures, suggesting an increase in larger high-mannose structures (**Figure 3d**). The predicted difference in oligomannose is consistent with reported EAS glycosylation; upstream phenylalanine (N-2) can decrease glycan processing and increase homogeneity.^17^ Structurally, W and Y are known to stabilize the chitobiose core through dispersion and increasing glycan accessibility for maturation; F lacks a dipole and therefore does not support similar maturation^97–101^. Both the EAS and selective aromatic stacking results are consistent with glycoimpact. We predict limited expected differential glycosylation following a W to Y substitution but a substantial decrease in processing following substitution from W or Y to F (**Figure 3f**, Supplementary **Figure 5**). The expanded sequon scaffolded by the IMR (Figure 2), glycoimpact substitution matrix (**Figure 3f**, Supplementary **Figure 5**), and glycosite-coupled residues (**Figure 4c-d**) together represent a portable and dynamic summary of our findings that can be easily applied to predict glycosylation following novel substitutions. We believe that the large number of both high glycoimpact and low structural impact substitutions (around 50% of AA have one such substitution) will be useful in understanding pathogenic variants, viral evolution, and glycoprotein engineering.

Glycoimpact can be further used to inform the mechanism of pathogenesis when disease variants are close to glycosites that impact glycosylation. We can examine known annotated-pathogenic variants and propose novel mechanisms of pathogenesis. Unannotated variants can be automatically labelled for likelihood of glycan modulation. Though this study does not deeply interrogate causal associations with glycan-modulated pathogenesis, we found several prior works implicating high-glycoimpact glycosite proximal variants as pathogenic, including one thorough study of oculocutaneous albinism implicating variant-proximal glycosylation as a causal pathogenic modulator. The highest glycoimpact relation with a high BLOSUM score, a Valine-Isoleucine substitution, is known to have a negligible protein structural impact while dramatically changing glycosylation. As expected, pathogenic variant PrP:V180I (adjacent to glycosite N181) is a causal mutation in Creutzfeldt-Jakob disease.^102^ Similarly, a glycosite-proximal V84I substitution in HIV-gp120 deactivates the virus; otherwise achieved by mutagenic glycosite-ablation.^103^ Similarly, tyrosinase:A355V (P14679) is a sufficient cause of oculocutaneous albinism.^52,53^ A355V is also a high glycoimpact variant and close to the function-critical N371 glycosite (<20Å; PDB:5M8N); tyrosinase glycosylation is critical for proper folding,^55,104^ N371 glycosite ablation results in decreased protein abundance and activity,^105^ and non-mutagenic post-expression tyrosinase deglycosylation also interrupts function.^57^ Using glycoimpact, we can propose A335V as a glycan-based mechanism of pathogenicity. Beyond tyrosinase, we identified thousands of variants across hundreds of diseases that may be similarly explained by aberrant glycosylation. While aberrant glycosylation can also be caused by environmental changes and proteostatic stress,^91,92^ glycoimpact opens a new an additional perspective on mechanisms by which aberrations in glycosylation may be transduced. There may be millions of high glycoimpact variants close to glycosites beyond the narrower scope of ClinVar annotation for which glycoimpact can suggest a possible pathogenic mechanism.

Just as amino acid substitution can assert pathogenic glycosylation on human proteins, similar substitutions can enable immune evasion for viruses. Consistent with glycoimpact predictions, we show that glycosite proximal proline and glutamine are associated with glycan complexity across HIV gp160. These results both validate our model and propose mechanisms for evolutionary control over the viral glycan shield.^14,28,106,107^ Across three SARS-CoV-2 Variants of Concern, we show that high glycoimpact variations close to glycosites produce IMR-consistent differential spike glycosylation. If IMR can predict differential glycosylation as viruses evolve, these predictions could help predict immune evasion and predict the viral evolutionary landscape for many viruses.^14,107,108^

Functional differential glycosylation is also critical in protein therapeutics. Glycoengineering for therapeutics is often costly, application-specific, and siloed from protein design. Glycans can be modified indirectly through genome editing of glycosyltransferases^109–111^ or media optimization,^112–114^ and such strategies can be guided by models of the biosynthetic pathways.^115–120^ Here, we examined AA-specific differential glycosylation events on antibodies. On the previously-characterized and differentially-glycosylated mouse IgG1 allotype (I299F), we see IMR-predictable glycosylation encoded directly into the glycoprotein primary sequence. Similarly, in glycosite-proximal and high glycoimpact mAb-Fc-variants, both fucosylation and fucose-modulated functional response (ADCC) correlated with GEE-IMR core-fucose preference. In the most extreme challenge, we show that novel Rituximab variants can be designed with IMR to deliberately and effectively change glycoprofiles. In these examples, IMR is a biologically^79–81^ and therapeutically^90,121–124^ useful method for connecting allotype, glycosylation, and effector function. Therefore, IMR present an opportunity to embed glycoengineering directly into the glycoprotein sequence and combine the currently separate paradigms of glycoengineering and protein engineering to a unified practice of glycoprotein engineering: considering the structure of both protein and glycan and their mutual influence.

Our direct results focus on human-expressed glycoproteins and global trends, rather than specific mechanisms such as aromatic chemistry, or chaperone proteins. While the specific glycoimpact trends may not extend to other organisms, the chemistry of carbohydrate-protein interactions suggests that glycoimpact is foundational and therefore exists in some form across glycosylated proteins. Additionally, these observations speak to glycosylation potential and will be most accurate in nascently-expressed proteins under limited proteostatic stress when glycogenes are expressed and substrates are available. We do not suggest these predictions to describe the only glycoprofile but rather the set of plausible glycans that may comprise a site-specific glycoprofile.

The evidence presented demonstrates the constraint of glycosylation by protein structure and the relevance of glycoimpact. These findings are corroborated by multiple distinct analyses and datasets with which protein structure appears to bound glycosylation. In all, protein structure appears to inform glycan biosynthesis. Predictability in glycosylation from protein structure will unlock a wealth of theoretical, exploratory, and corroborative analysis making it inexpensive and easy to leverage glycobiological insights in fields throughout biology.

## Acknowledgements

Thank you to Terry Platt who inspired me with tales of the *Trp* operon^131–135^ and in doing so planted the seed of thought that substrate-limitation is not mutually exclusive with templated biosynthesis. Thanks also to Lorenzo Casalino, Christian Seitz, and Rommie Amaro for their early insights into the project in the context of influenza glycoprotein structure. We would like to acknowledge the invaluable contribution of Dr. Jasminka Krištić, Prof. Dr. Grant Morahan and Prof. Dr. Falk Nimmerjahn to the studies of IgG N-glycome that are referred to in this paper. EA is supported by ANID (DOCTORADO BECAS CHILE/2018 - 72190270), the Fulbright Chile Commission, and the Siebel Scholar Foundation. This work was supported by NIGMS (R35 GM119850, NEL) and the Novo Nordisk Foundation (NNF20SA0066621, NEL).

## Methods

### Enrichment of glycan-protein site-matched data to generate the Protein-Glycan Enriched Structure Dataset (PGD)

Starting from site-specific glycosylation events, we extended the annotation of each glycosylation site and glycan to include detailed site-specific protein structural annotation and recorded the number of times each substructure (defined below) appeared in each glycan. Only human glycoproteins were analyzed. The final database includes 111 human glycoproteins (98 glycoproteins with N-glycosylation sites and 38 glycoproteins with O-glycosylation sites) 306 glycosylation sites and 4,263 glycans (3,563 N- and 700 O-linked glycans). We initially used site-specific glycosylation events documented in UniCarbKB (original deprecated, latest version archived at data.glygen.org/GLY_000040) and in the January 2022 release of GlyConnect^37^ (glyconnect.expasy.org/)).^41^ Later and current work was informed by glycosylation events documented in GlyConnect^136^ with supplemental information from GlyGen.^137^ Known glycosylation events from the UniCarbKB and GlyConnect were used to inform much of the core analysis. Glycomes and glycoproteomes in UniCarbKB were collected from GlycosuiteDB,^138^ EUROCarbDB,^139^ and expanded meta-analysis data.^27^ GlyConnect was built on the fall 2017 release of UniCarbKB and spans glycomes and glycoproteomes predominantly (70%) curated from experiments distinct from UniCarbKB.

The protein structure annotation was done using the Structural Systems Biology (ssbio) package in python.^39^ The package uses several tools to perform a variety of annotations. For each human protein, empirical and homology modeled structures were collected from the Protein Data Bank (PDB)^140^ and SWISMOD,^141^ respectively. Proteins without existing models were modelled using I-TASSER.^142^ Protein structures and chemistry close to the glycosylation sites were annotated multiple software packages through ssbio: sequence properties (EMBOS:*pepstats*),^143^ sequence alignment (EMBOS:*needle*),^143^ secondary structure (DSSP,^144^ SCRATCH::SSpro^145^ and SCRATCH::SSpro8), solvent accessibility (DSSP and FreeSASA), and residue depth (MSMS). Additional amino acid aggregate features were calculated using R::seqinr. Spatial proximity was defined using “min-distance” between two amino acids; the minimum distance between any pair of atoms spanning the amino acids.

Glycan structures were used to represent shared structures between specific observations. Several authors have relied on the representation of glycans as graphs starting with early computer encodings such as KCF^146^ or GlycoCT^147^ and all the way to latest glycan language models^148^. Grounding our work in this trend, we consider a substructure as a subgraph of the full graph that represents a whole structure. Substructures were annotated using a combination of glypy^149^ and GlyCompare^38^ for structure parsing and comparison respectively. All glycan *substructures*, a connected subset of monosaccharides with and without linkage information, were extracted from each glycan, merged to make a superset of substructures, then mapped to each glycan. Thus, resulting in a mapping from every glycan in the input database to shared substructures. We define *motifs* as notable glycan substructures.

### Software and packages

Protein structure analysis was performed in Python v2.7.15 using ssbio v0.9.9.8 to retrieve and calculate: existing empirical and homology models from PDB and SWISSMOD (PDBe SIFTS),^150^ *de novo* homology models (I-TASSER v5.1), sequence properties (EMBOS v6.6.0.0 pepstats), sequence alignment(EMBOS v6.6.0.0 needle), secondary structure (DSSP v3.0.0, SCRATCHv1.1::SSpro and SCRATCHv1.1::SSpro8), solvent accessibility (DSSPv3.0.0 and FreeSASAv2.0.2), and residue depth (MSMSv2.2.6.1). Additional amino acid aggregate features were calculated using R::seqinr.

Statistical analysis was performed in R v3.6.1. R::entropy v1.2.1 was used for entropy, Kullback-Leibler divergence and other information-theoretic calculations. Generalized Estimating Equations (GEE) were fit using R::geepack v1.3.1. Gaussian Mixture Models were used to z-score normalize the glycoimpact using R::mixtools v1.1.0. BLOSUM and PAM substitution matrixes were accessed from R::Biostrings v2.52.

### Probability event space, information gain and conditional probability

Here we define an event (a row in our enriched glycosylation-glycosite database) as “the observation of a glycan at a glycosylation site in an experiment.” If two separate experiments in the input database both report the same glycan at the same site on the same protein, we consider that event to have occurred twice. Within each event, we ask if the glycan structure random variable (the presence or absence of a specific glycan substructure) is present or absent in the observed glycosylation event and if the protein structure random variable (a proximal amino acid, a secondary structure or another discrete protein structural feature). A Fisher exact test (R::base::fisher.test) was used to estimate the odds ratio (OR) and significance (p) of each intra-molecular relation (IMR). P-values were corrected for False Discovery Rate (FDR, q) permitting 10% false discovery (q<0.1); a common threshold for systems-level analyses and distinct from p-values. Conditional probability was calculated by dividing joint probability by the marginal probability of protein and glycan structure presence. Kullback-Leibler divergence (KLd, R::entropy::KL.Dirichlet, pseudo count=1/6) was calculated by comparing the conditional probability distribution to the marginal probability distributions. In summary, we quantify an individual IMR as the Fischer exact test odds ratio (OR), significance (p), that were corrected for by the false discovery rate (FDR), which were later analyzed by clustering.

### Quantitative characterization of Intra-Molecular Relations (IMR) using Generalized Estimation Equations (GEE)

To characterize the IMR in the PGD while controlling for protein-specific confounding effects and handle nonlinear relations we used a population-averaging approach; logistic Generalized Estimating Equations (GEE) with glycoprotein identity as the cluster identifier.^151^ We used an exchangeable correlation structure to describe and balance the in-protein similarity. Models were fit to predict glycan substructure binary (presence or absence) from either z-score-normalized continuous or binary (presence or absence) protein structures. For each model, the data from PGD was isolated for one glycan-type (N-glycan or O-glycan), one glycan substructure and one protein structure. Incomplete observations (events/rows) were removed and then several checks on each data-slice were run to minimize overfitting. Glycan substructures were excluded from modelling if standard deviation was less than 1e-6 or if there were fewer than 5 observations of the structure within the pertinent data-slice. Discrete protein structure features were excluded if there were fewer than 4 observations within the data-slice. Models were excluded if there were fewer than 4 instances in any cell (of the 2×2 absence/occurrence matrix) or if the chi-squared expected value of any cell was less than or equal to 5. Observations were weighted by the reciprocal-count of the corresponding label type to balance label contributions to the model and scaled by exponentiated c-score to maximize the contribution of high-quality protein structure models (*w*_*i*_ = 2^*C*^ / *n*); *c* is the c-score given by I-TASSER and n is the number of times a structure is present (1) or absent (0). Models with |log(OR)|> 50 were excluded as likely overfit. Quasi-likelihood under independent model criterion (QIC) and the Wald tests were used to evaluate the significance and magnitude of the estimated IMR. We also ran this analysis using publication identifiers as a group/cluster identity variable to account for researcher and group biases; this produced similar results likely because protein identity is strongly correlated with the publications in which the proteins appear.

### Calculating glycoimpact from IMR and populating a BLAMO matrix

Glycoimpact is the z-score normalized Euclidean distance between the significant log(OR) for all motifs associated with a protein structure; the null distribution of Euclidean distances was determined using Gaussian mixture models (Supplementary **Figure 5**) Glycoimpact is calculated for every pair of AAs as the Euclidean distance between significant and substantial log odds ratios for each AA; the Euclidean distance between expected glycoprofiles for each AA. The substantial (log(OR)>X) and significant (FDR<Y) log(OR) values are retained while insignificant or unsubstantial log(OR) values are set to zero. The resulting matrix describes the expected glycoimpact due to each AA-substitution, termed the BLAMO XY matrix where X and Y denote the log(OR) and FDR thresholds respectively.

Glycoimpact values from a BLAMO XY matrix can then be z-score normalized to a Gaussian Mixture Model^152^ estimated null distribution. We use z=2.5 as a heuristic but stringent cutoff between “impactful” (z>2.5) and “null” (z<2.5) substitutions.

### Comparison of SNP pathogenicity scores with glycoimpact

Functional prediction rank normalized scores were obtained from dbNSFP (v3.2) for the following 27 tools: SIFT, PolyPhen-2 HDIV, PolyPhen-2 HVAR, GERP++, MutationTaster, Mutation Assessor, FATHMM, LRT, SiPhy, 2x PhyloP, MetaSVM, MetaLR, CADD, VEST3, PROVEAN, 4× fitCons scores, fathmm-MKL, DANN, 2× phastCons, GenoCanyon, Eigen and Eigen-PC.^49^ Variants were excluded from the analysis if they had more than 3 missing functional score predictions, did not result in an amino acid change, or not on proteins that had known glycosylation sites.

Assignments of “prediction-type” and “structure-usage” (**Figure 5b**) were adapted from classifications provided by dbNSFP.^49^

### Estimation and Analysis of Evolutionary Couplings (EC)

For EVCouplings calculation, hits composed of more than 50% gaps were filtered from the alignment, and sequences with homologs more than 80% identical were down-weighted to compute Neff, the effective number of sequences.^48^ ECs were calculated using pseudo-likelihood maximization,^153,154^ as implemented previously.^155^ The λ_J_ term was scaled by the number of amino acids minus one times the number of sites in the model minus one. Pre- and post-processing was performed using the EVCouplings Python package.^156^

High-ranking EC events are generally considered those ranking less than *L* a measure of sequence length where only residues with fewer than 30% gaps were counted.^48^ Specifically, the gap threshold (minimum column coverage parameter) was set to 70%. Therefore, residue positions with more than 30% gaps disregarded. We also used a fragment filter of 70%, meaning that individual sequences had to have 30% non-gap characters in each row as well. We explored multiple high-rank thresholds between *L*/5 and 3*L*. To explore the increased coupling with glycosylation sites, we examined couplings between each amino acid with glycosites (GN), asparagines (N) and any amino acid (AA). We compared the number of high-ranking coupling events, the distributions of EC probabilities and the relative numbers of high and low-ranking ECs for each group with various amino acids at relative positions N+/-6. Distributions were compared with a one-sided Wilcoxon test and high/low-ranking counts were compared with hypergeometric enrichment. The hypergeometric enrichment of glycosite-coupling was performed at multiple high-rank thresholds (*L/3, L/2, L, 2L, 3L*) and p-values were pooled for each amino acid at each relative position across ranks using Fisher’s method. Finally, the pooled p-values were corrected for multiple tests using the Benjamini-Hochberg method.

To examine larger structures in ECs, we used EC rank to mask extended sequons (N+/-6) then clustered the sequons and extracted motifs. For each sequon, the residues were retained if the residue-glycosite coupling rank was less than *L4*. The extended and masked sequons were distinguished using a hamming distance (DECIPHERv2.18.1)^157^ then clustered using agglomerative hierarchical clustering (factoextra::hcut v1.0.7). Motif logos were generated using custom-scaled position-specific scoring matrices^158^ reflecting the cumulative rank of amino acids at each glycosite relative position. Specifically, the aggregate score, *S*, for each amino acid, *a*, at each position, *p*, was aggregated over EC score ranks, *r*, within each extended-masked sequons, *s*, in a cluster, *c*, such that *s*_*a,p*_ = *Log*_10_ ∑_*S* ∈ *c*_ *L*/*r*)

### Mouse breeding and Samples

The Collaborative Cross (CC) recombinant inbred mouse strains (N = 333, 95 strains, age 20-117 weeks) were produced by Geniad Pty Ltd and housed at Animal Resources Centre (Murdoch, WA, Australia).^159^ The CC strains were genotyped using the MegaMUGA platform (GeneSeek; Lincoln, NE). C57BL/6 (N = 10) and BALB/c mice (N = 10), sex- and age-matched (10 weeks old, 1:1 male:female) were obtained from Elevage Janvier (Le Genest-Saint-Isle, France).^89^ The studies received appropriate ethics approvals from the Animal Ethics Committee of the Animal Resources Centre^78^ and the Ethical Committee of the District Government of Lower Franconia.^89^

### Liquid Chromatography – Mass Spectrometry (LC-MS), Normalization and Statistical Analysis of Mouse Fc-linked IgG N-glycopeptides

Immunoglobulin G was isolated from 100-500 μl of mouse serum on 96-well Protein G monolithic plates (BIA Separations) as described previously.^78,88^ LC-MS analysis of tryptic Fc-glycopeptides was performed as described in.^78,88^ In brief, approximately 10–20 μg of isolated IgG was digested with 200 ng trypsin (Worthington, USA). The resulting glycopeptides were purified by reverse-phase solid phase extraction using Chromabond C18ec beads (Marcherey-Nagel, Germany) as described in.^88^ Tryptic digests were analyzed on a nanoACQUITY UPLC system (Waters, USA) coupled to a Compact mass spectrometer (Bruker Daltonics, Germany). Peak areas were calculated by summing areas for doubly and triply charged ions determined with LaCyTools v 1.0.1 b.7 software^160^ and normalized to the total integrated area per IgG subclass.

Batch correction was performed on the log-transformed values using the ComBat method (R package “sva”) to remove possible experimental variations due to LC-MS analysis having been performed on several 96-well plates within each cohort. Derived glycosylation traits describing relative abundance of N-glycans sharing specific structural features (agalactosylated, galactosylated, sialylated, monogalactosylated, digalactosylated, monosialylated, disialylated structures, structures with bisecting GlcNAc) were calculated in a subclass-specific manner.^89^ Statistical analysis and data visualization were performed using R programming language v 4.0.3.

### Glycoproteomics for SARS-CoV-2 Spike glycoproteins

His-tagged recombinant SARS-CoV-2 Spike S1 protein, expressed in HEK293 cells were purchased from Sino Biologics (Wayne, PA). Lyophilized glycoproteins corresponding to the original 2019 strain (40591-V08H), the Gamma variant (40591-V08H14 with L18F, T20N, P26S, D138Y, R190S, K417T, E484K, N501Y, D614G and H655Y mutations), and the Delta variant (40591-V08H23 with T19R, G142D, E156G, 157-158 deletion, L452R, T478K, D614G and P681R mutations). Samples were analyzed using DeGlyPHER.^77^

Briefly, glycoproteins were digested with Proteinase K (30 min or 4 h) or trypsin, sequentially deglycosylated with Endo H (creating residual mass signature of +203 Da) to signify high mannose/hybrid glycans, and then with PNGase F in H2^18^O (creating residual mass signature of +3 Da) to signify the remnant complex glycans on any sequon (NXS|T, where X is any amino acid except P) asparagine. Unoccupied sequons will have no additional signature mass. Analysis of samples was done on a Q Exactive HF-X mass spectrometer (Thermo), injecting directly onto a 25 cm, 100 μm ID column packed with BEH 1.7 μm C18 resin (Waters). Liquid chromatography separation was achieved at a flow rate of 300 nL min−1 on an EASY-nLC 1200 (Thermo). Buffers A and B were 0.1% formic acid in 5 and 80% acetonitrile, respectively. The gradient used was 1−25% B over 160 min, an increase to 40% B over 40 min, an increase to 90% B over another 10 and 30 min at 90% B for a total run time of 240 min. The column was re-equilibrated with solution A before injecting sample. Peptides eluting from the tip of the column were nanosprayed directly into the mass spectrometer by application of 2.8 kV at the back of the column. The mass spectrometer was operated in a data-dependent mode. Full MS1 scans were collected in the Orbitrap at 120000 resolution. The 10 most abundant ions per scan were selected for HCD MS/MS at 25 NCE. Dynamic exclusion was enabled with a 10 s duration and +1 ions were excluded. Peptides were identified with Integrated Proteomics Pipeline (IP2, Bruker Scientific LLC). Tandem mass spectra were extracted from raw files using RawConverter^161^ and searched with ProLuCID^162^ against a database comprising UniProt reviewed proteome for Homo sapiens (UP000005640), including additional UniProt amino acid sequences for Endo H (P04067), PNGase F (Q9XBM8), and Proteinase K (P06873), the amino acid sequences for the SARS-CoV-2 S1 subunits, and a list of general protein contaminants. The search included no protease specificity (all fully tryptic and semitryptic peptide candidates when treated with trypsin). Carbamidomethylation (+57.02146 C) was used as a static modification. Deamidation in the presence of H_2_^18^O (+2.988261 N), GlcNAc (+203.079373 N), oxidation (+15.994915 M), and N-terminal pyroglutamate formation (−17.026549 Q) were used as differential modifications. Data were searched with 50 ppm parent mass tolerance and 50 ppm fragment mass tolerance. Identified proteins were filtered using DTASelect2^163^ while using a target-decoy database search strategy to limit the false discovery rate to 1%, at the spectrum level.^164^ At least one peptide per protein and no tryptic end (or one tryptic end when treated with trypsin) per peptide were necessary, and precursor delta mass cutoff was fixed at 10 ppm. Statistical models for peptide mass modification (modstat) were applied (trypstat was additionally applied for trypsin-treated samples). Semi-quantitative label-free analysis was performed based on the precursor peak area, with a 10 ppm parent mass tolerance and 0.1 min retention time tolerance, Census2.^165^ “Match between runs” was used to find missing peptides between runs. GlycoMSQuant (v.1.4.1, https://github.com/proteomicsyates/GlycoMSQuant)^77^ was used, summing precursor peak areas across the 3 conditions – 30 min and 4 h Proteinase K, and trypsin, discarded peptides without glycosites, and discarded misidentified peptides when N-glycan remnant mass modifications were localized to non-glycosite asparagines and corrected/fixed N-glycan mis-localization where appropriate, to finally calculate proportions of 3 glycosylation states at each glycosite, with +/-SEM (standard error of mean). Pairwise-statistical comparison between variants was performed using Mann-Whitney U Test.

The entire DeGlyPHER pipeline and measurement were run twice to collect two independent technical replicates. Using the Fisher’s method for pooling independent p-values, we combined the Mann-Whitney U-derived p-values for each glycosite comparison. Pooled p-values were adjusted for multiple-testing using FDR. The N17 glycosite was inconsistently cleaved with the signal peptide precluding stable measurements needed for robust comparison.

### Estimation of glycomotif preference from IMR

Here, we define glycomotif (e.g., core-fucose) preference. Briefly, this is the preference for a glycomotif following a substitution compared to the wildtype (WT) and a close (within one monosaccharide) precursor of the glycomotif of interest. When the IMR relating the lost (WT, x-axis) and gained (variant, y-axis) amino acids association (IMR) to the glycomotif and glycomotif precursor, the variant preference can be calculated as the distance between the glycomotif and glycomotif-precursor points; distance is the component perpendicular to the line of equity (y=x). Supplementary **Figure 15** demonstrates this calculation visually.

Glycomotif preference can be described in more precise terms. Given a substitution where *X* and *Y* are WT and mutant amino acids respectively at an amino acid residue at index,, we may describe the substitution as *XIY*. When *I* is glycosite-proximal, we may describe a substitution, *XIY*, as a point,, with a pair of IMR (ρ) relating each amino acid to a single glycomotif, *p*_*XIY*_ =(*ρ*_*X*_, *ρ*_*Y*_). The IMR-pair, *p*_*XIY*_ =(*ρ*_*X*_, *ρ*_*Y*_), may also be described as a vector terminating on that point, 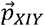. We may also represent the line of equality (*y*= *x*) by the normalized vector 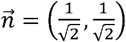. Here, the glycoimpact is the minimum distance, 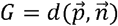 . To calculate the component of 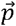 parallel to 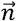, we project 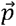 onto 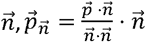. Therefore, 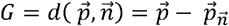

With this definition of *G*, we can define *G* _*i*_ and *G* _*C*_ as the glycoimpact corresponding to the glycomotif of interest and the contract glycomotif respectively. Then, the relative preference by first calculating the perpendicular component of the difference in glycoimpact vectors projected onto 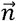, such that 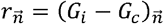. Then *R* is the *l* ^2^-norm of the vector difference, 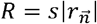. For 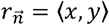, we define the sign,*s*, as positive if *y*> *x* and negative if *x*> *y*. The sign is not defined if *x*= *y*.

### Glycoprofiling of 25c4 monoclonal antibody using capillary electrophoresis

As was recently described,^166^ 54 Fc variants of 25c4 were cloned from the REFORM Fc variant panel.^90^ The REFORM plasmid library^90^ is comprised of golden gate cloning plasmids^167^ with Bsal restriction sites flanking distinct antibody domains and a furin 2A cleavage site to enable self-cleavage and successful assembly of a complete antibody from a single open reading frame.^168^ Variable Light chains containing REFORM variants and 25c4 Variable Heavy chains were simultaneously transfected at a 1:1 ratio in CHO cells, purified using a Protein A chromatography resin, then dialyzed and concentrated with Phosphate Saline Buffer (PBS).

Fc glycosylation was then measured on purified 25c4 antibodies. Purified antibodies were incubated with magnetic G protein beads (Millipore) and separated from Fab fragments following enzymatic digestion (IdeZ, NEB). Fc glycans were released and labelled using the GlycanAssure ATPS kit (Thermo Fisher Scientific) then separated using 3500xL Genetic Analyzer (Thermo Fisher Scientific). As previously described,^169^ retention times (RT) were matched to glycan standards using Glycan Acquisition Software Version 3500 v1.0.3 and Glycan Analysis Software v1.1. Abundance (area under peaks) was normalized to total area per sample to calculate relative abundance of each glycan. Non-uniquely determined glycans were excluded (where the difference in RT was below the detection threshold).

### Expression, purification and glycoprofiling of Rituximab variants using LC-MS-backed UPLC

Rituximab titers were determined in triplicate on an Octet® Red96 biolayer interferometry (BLI) instrument. Binding to ProA biosensors was recorded for 120 s at 30 °C. Binding rates were converted to concentrations based on a standard curve generated using wildtype Rituximab, produced and purified in-house.

The Rituximab variants were purified by affinity chromatography using a 1-mL MAb Select Sure column (Cytiva) mounted on an Äkta Pure instrument. Equilibration and washing steps were performed using 20 mM sodium phosphate, 0.15 M NaCl, pH 7.2. The antibody was eluted with 0.1M Sodium citrate, pH 3. Elution fractions were neutralized with 0.2 V of 1 M Tris, pH 9. Next, the protein solutions were desalted using 5-mL Zeba™ Spin desalting columns (7K MWCO, Thermo Fisher) and dPBS as eluent. Finally, the desalted solutions were concentrated on 4-mL Amicon centrifugal filter units (50K MWCO, Millipore) aiming for a concentration of approximately 0.5 mg/mL. The final concentrations were determined by measuring absorbance at 280 nm on a Nanodrop 2000 spectrophotometer using an extinction coefficient of 1.46 (mg/mL)^-1^cm^-1^.

Purified and concentrated protein extract were fluorescently labelled (N-glycan labeling using the GlycoWorks RapiFluor-MS N-Glycan Kit, Waters, Milford, MA). Fluorescent glycans were stratified and measured using a HILIC-FLR with ACQUITY UPLC Glycan BEH Amide column (2.1 x 150 mm, 1.7 µm, Waters, Milford, MA) mounted on an Ultimate 3000 UPLC system and a Fusion Orbitrap mass spectrometer (Thermo Scientific). Acetonitrile (100%) and ammonium formate (50 mM, pH 4.4) were used as mobile phases.

