## Supplemental for "Protein structure, a genetic constraint on glycosylation"

### Supplementary Materials: Protein structure, a genetic encoding for glycosylation

#### Supplemental Results

##### PGD glycosites are generalizable and represent all glycosite structures

We first verified that the glycosite-structure annotation represents typical protein structure variation at glycosites. With the structure-annotated glycosites, we performed a dimensionality reduction (Factor Analysis for Mixed Data (FAMD), Supplementary **Figure** 2a-b). We then projected a test set of all Uniprot-annotated glycosites in the human secretome[42] into the reduced space (these glycosites were not in the FAMD training set). Using a multivariate Gaussian, we determined the probability that each test-set glycosite was within the distribution of input glycosite variation. After a False Discovery Rate (FDR) correction,[170] we found no outlying glycosite structures (FDR<0.1, Supplementary **Figure** 2a-c), indicating that the PGD glycosites are representative of the broader space of glycosites. Thus, PGD can provide generalizable conclusions about protein-glycan associations.

##### Glycan and protein structures are not independent

We examined the associations (termed "intramolecular relations (**IMR**)") between glycan substructures (e.g., tri-mannose) and glycosite-proximal protein features (e.g., proximal tyrosine) within PGD (Supplementary **Figure** 3a). IMR are first computed using the Fisher exact test odds ratio (OR). To examine asymmetries in IMR, we explored non-independence (NI, the absolute difference between conditional and marginal probabilities) and Kullback–Leibler divergence (KLd) between glycan and protein presence. Specifically, we examined conditional and marginal distributions of protein structure when the glycan structure was fixed, present (G=1) or absent (G=0) and glycan structure when the protein structure was fixed, present (P=1) or absent (P=0). Due to a large sample size, most mean differences were trivially significant, and thus, only effect size is reported.

Of 259,114 potential substructure-IMR, we found 50,842 relationships that were substantial (|Fisher-OR|>0.1) and significant (Fisher-FDR<0.1). Of 26,404 motif-IMR (numerically and biosynthetically non-redundant substructure-IMR),[38] 10,111 were substantial and significant. Of the 10,111 selected motif-IMR, there were 9,296 positive and 815 negative correlations. The most significant IMR included correlations with glycosite-proximal (within 6Å) alanine and cysteine residues, and anticorrelations with proximal arginine and valine residues (Supplementary **Figure** 3**b**). We further analyzed the NI and KLd when either protein structure or glycan structure was known (Supplementary **Figure** 3**c**). KLd was low for significant IMR (Fisher exact test; FDR<0.1) when protein structure or glycan structures were not present (Mean $KLd_{P=0}$ = 0.0038, Mean $KLd_{G=0}$ = 0.0054). KLd increased ~10-fold when protein structure was present (Mean $KLd_{P=1}$ = 0.032) and ~25-fold when glycan structure was present (Mean $KLd_{G=1}$ = 0.138). Overall, the presence of glycan structure provides the most information about protein structures, whereas the protein structure provides lesser, but substantial information about glycan structure.

We examined motif-IMR by estimating the conditional probability between protein and glycan structures. Conditional probability diverged significantly (Fisher exact test; FDR < 0.1) from corresponding marginal probabilities, indicating non-independence (Supplementary **Figure** 3**d-e**). Conditional glycan probabilities (Supplementary **Figure** 3**d**) and conditional protein structure probabilities (Supplementary **Figure** 3**e**) show symmetric (Loess estimation) difference from respective marginal probabilities suggesting no global bias for either condition. We also stratified the glycan-protein structure non-independence by glycan motif size (number of monosaccharides, Supplementary **Figure** 3**f**, Supplementary **Table** 4). For monomeric glycan motifs, protein structure was 2.3-fold less determined by glycan structure (Mean $NI_{P|G}$ = 0.091, $NI_{G|P}$ =0.040). As motif size increases, the disparity in non-independence grows to 34.2-fold increase at 21-mers (Mean $NI_{G|P}$ = 0.038, $NI_{P|G}$ = 0.10). These results suggest larger glycans are less determined by protein structures but better inform protein structure. Indeed, glycosylation influences protein folding.[171,172] The decrease in non-independence in larger glycans is consistent with our bounded biosynthesis hypothesis. While proteins may request glycan structures, more mature and larger glycan structures are less likely to appear when requested due to enzyme and substrate limitations; $NI_{G|P}$ may not diminish in larger oligomannose which may explain the NI stability for motifs up to 11-mers (the largest oligomannose). The moderate-high non-independence of glycan structure on protein structure is consistent with our premise that glycosylation is not template-free biosynthesis.

#### Amino acid changes can predict glycan structural changes

We found that many AA-substitutions can have substantial "glycoimpact" (expected difference in glycosylation) on glycan biosynthesis; exemplified by the phenylalanine-tryptophan substitution (**Figure** 3**d**). An upstream phenylalanine is associated (>3-fold) with core fucosylated tri- and bi-antennary structures with variable galactosylation. Upstream tryptophan is marginally associated (<2-fold) with core-fucosylated biantennary structures too but more associated (>2-fold) with tetraantennary structure. Thus, a phenylalanine-tryptophan substitution could impact branch number and core-fucosylation.

Interestingly, upstream phenylalanine is associated (>3-fold) with a Man7 substructure but anti-correlated with a Man6 substructure (>2-fold) suggesting that upstream phenylalanine, in some contexts, prefers larger oligomannosidic structures. At the structural-level, proximal phenylalanine and tryptophan show related effects. Spatially proximal phenylalanine is correlated (>10-fold) with an increase in sialylation on triantennary core-fucosylated structures while tryptophan is correlated with distal fucosylation (>2-fold) (**Figure** 3**e**). These predictions suggest that tryptophan-phenylalanine substitution has a large impact on glycoprofiles.

#### High glycoimpact amino acids show increased evolutionary coupling with N-glycosylation sites

If glycosite-proximal amino acids influence functional glycosylation, there should be evolutionary pressure imposed beyond the classical NX[S/T] N-glycan sequon. Thus, we calculated evolutionary coupling (EC) scores from functional-domain alignments of 2,005 glycoproteins—EC are derived from a global probability model and accurately predict structural and functional relationships between amino acids from co-evolution.[48] We examined AA pairs from the top *n* ranked EC scores using multiple score cutoffs from n=L/5 to n=4L—sequence length (L) across a multiple sequence alignment counting residues with fewer than 30% gaps. We first examined the number of high-ranking EC (above rank threshold) between any amino acid (X) with N-glycosylation sites (GN). At multiple rank thresholds (n=L/5 to n=L/3), we found significantly more high-ranking glycosite-coupled EC (GN) than Asn-coupled (N, p<0.029, one-sided Wilcoxon-test) or all background EC (X, p<0.0013, one-sided Wilcoxon-test, Supplementary **Figure** 7a).

We next compared GN, N, and X couplings with upstream (N-terminal (N-i)) or downstream (C-terminal (N+i)) amino acids. We measured glycosite coupling with another position-specific amino acid by evaluating a continuous and a discrete coupling metric. Using continuous coupling probabilities, we examined increased GN-coupling probability relative to N- or X-coupling at a given rank threshold (one-sided Wilcoxon test, Supplementary **Figure** 7**b**). Using the rank threshold dictated top-ranking vs null EC distinction, we examined the increased proportion of high-ranking GN-coupled events relative to comparable N- or X-coupling (Supplementary **Figure** 7c)—increased proportion was measured by hypergeometric enrichment at multiple rank thresholds then pooled across thresholds using Fisher's method and corrected for multiple-testing (FDR) (see **Methods**). As expected, given the N-glycan sequon, serine and threonine at N+2 have significantly higher coupling probability with glycosites than other Asn (N) or background (X) (One-sided Wilcoxon p<0.05, Supplementary **Figure** 7b). Of top-ranking EC, at multiple thresholds, those involving N+2 serine and threonine are more likely to couple with glycosites than other Asn (N) or background (X) (pooled hypergeometric FDR<0.1, **Figure** 4**c**). We found several additional position-specific glycosite-coupled residues, including phenylalanine at N-2 (hypergeometric $p<0.005$, Supplementary **Figure** 7c; pooled hypergeometric FDR < 0.1, **Figure** 4**c)** and tyrosine at N-1 (pooled hypergeometric FDR < 0.1, **Figure 4c**). These findings are consistent with previous observations of the enhanced aromatic sequon.[17]

Examining position-specific glycosite couplings (hypergeometric enrichment for high-rank EC pooled across rank-thresholds, **Figure** 4**c,** Supplementary **Figure** 7c), we found 13 of 20 AA have at least one significant (FDR<0.1) increase in co-occurrence with glycosites over other asparagines (N, red-square) or any AA (X, black triangle). Seven of ten AA strongly implicated in upstream glycosite interactions (those visible in **Figure** 2**c, "Asn-5 N-term"**) show enriched coupling with glycosites; specifically, A (N-1,2,4,6), D (N-2), F (N-2), I (N-2), K (N-2), L (N-1,3,5), and S (N-2). Several glycosite-coupling events are enriched over either N or X but not both. When glycosite coupling probabilities were clustered (**Figure** 4**d**), glycosite-flanking amino acid motifs begin to resolve. Glycosites (N+/-6) were annotated with high-ranking EC scores (only rank n<4L were retained), then motifs were clustered, and motifs were constructed for 5 motif-clusters (**Figure** 4**d**) and m25 motif-clusters (Supplementary **Figure** 8, see **Methods**). The N+2 aspartic acid enriched in the univariate analysis (**Figure** 4**c**) co-occurs with an N-2 K (**Figure** 4**d,** motif 1). Alternatively, E is more likely to co-occur with other E residues (N -4, +1, and +3) with an N+2 T sequon (**Figure** 4**d,** motif 4). These couplings, reflective of evolutionary pressures, surrounding the glycosylation sites suggest a dramatic expansion of the N-glycosylation site structure.

Finally, we aligned[15] glycosites permitting a tetraantennary N-glycan lacking fucose or sialic acids (**Figure** 4**b**). We examined the glycosite alignment for consistency with high-influence AA (**Figure 2c**) and those significantly coupled with glycosylation sites (**Figure** 4**c**). Of 20 glycosite-flanking AA, 16 show consistency between the first or second most common AA and either the high-influence or glycosite coupled residues. We tested binomial enrichment from 20 possible AA where 10 high-influence upstream AA may appear at any of 10 upstream positions, or 8 high-influence downstream AA may appear at any of 9 downstream positions (**Figure 2c**) in the primary glycosite consensus sequence (PWQAKVVSRHNLTQGATLLNE, N+/-10, **Figure** 4**b**). We observed 5 high-influence residues appearing upstream (S, K, A, Q, W; binomial $N=10$, $p=10/20$, $Pr(X>5)=0.377$) and the 6 high-influence AA appearing downstream (T, Q, G, T, L, L; binomial $N=9$, $p=8/20$, $Pr(X>6)=0.025$) Glycosite-coupled residues—39 coupled AA / 19 positions = 2.05 residues per position (**Figure** 4**c**)—in the primary consensus sequence are enriched upstream in the glycosite alignment (P, A, V, S, H; binomial $N=10$, $p=2.05/10$, $Pr(X>5)=0.00724$). At nearly every glycosite-flanking residue (N+/-10) there is consistency between these three analyses, further corroborating that protein structure constrains glycosylation.

##### Silent substitutions are pathogenic without influencing protein structure

As expected, substitutions that disrupt protein structure (low BLOSUM score) are enriched among pathogenic ClinVar variants. Conversely, variants with high BLOSUM scores (non-disruptive to protein structure) are enriched among benign ClinVar variants (Supplementary **Figure** 17A). While many pathogenicity predictions conform to this pathogenic-BLOSUM association, FATHMM[173–175] shows only a muted form of the trend suggesting that there are other, possibly underutilized features influencing pathogenicity beyond strict protein structure disruption (Supplementary **Figure** 17B). Towards exploring pathogenicity beyond protein structure disruption, we stratified pathogenic substitutions (across BLOSUM scores and marginalized on clinical significance) by glycosite proximity (within 10A sphere, minimum distance) and found that the pathogenic-BLOSUM association is sensitive to glycosyite proximity. Specifically, a high-BLOSUM (non-disruptive) pathogenic variant is nearly 10% more likely to occur close to a glycosite (Supplementary **Figure** 17C-D). The enrichment of protein structure-ambivalent, yet pathogenic variants close to glycosites highlights a yet unexplained and likely overlooked phenomenon in genetic pathogenicity.

#### Supplemental Discussion

##### Glycogenomics in mouse IgG

Immunoglobulin (Ig) G is one of the most abundant N-glycosylated proteins in the blood plasma of mammalian species. Its N-glycome is well-characterized in humans[176] and mice,[78,87] the latter being an important model organism for immunological studies. Each human and murine immunoglobulin G heavy chain possesses a conservative N-glycosylation site in the CH2 domain of the fragment crystallizable (Fc) domain. Also, up to 20% of IgG molecules are N-glycosylated in the variable antigen-binding (Fab) region. IgG N-glycome mainly consists of complex type biantennary N-glycans. The glycan structures found on IgG exhibit typical conserved N-glycan core, with modifications that may or may not be present, such as core-fucose, bisecting N-acetylglycosamine (GlcNAc), and optional decoration of one or both antennae with galactose residues, that can be further decorated with sialic acid residues (**Figure** 7**c**). IgG N-glycosylation is a complex trait defined by both environment and genetics.[177,178] Regulation of IgG N-glycosylation is realized through a complex network of interactions between the glycosylated protein, glycosyltransferase and glycosidase enzymes, Golgi milieu and composition, and is rather difficult to study due to the absence of a direct constraint for glycan biosynthesis and the number of enzymes involved.[3]

However, there is evidence that one of the factors affecting IgG N-glycome might be the amino acid composition of the IgG heavy chain. For instance, a point mutation Tyr407 introduced to the CH3 domain of human IgG led to increased galactosylation, sialylation and branching of N-glycans attached to Asn297 in single heavy-light chain pairs expressed in cell cultures.[179] In the other experiment, human IgG3 variants with amino acid residues interacting with the Fc-linked N-glycan mutated to alanine were expressed in CHO cells[12]. The analysis showed that substitutions FA241, FA243, VA264, DA265, and RA301 led to increased abundance of digalactosylated and sialylated structures, while the YA296 substitution, on the contrary, resulted in IgG3 almost devoid of N-glycan structures with terminal sialylation and galactosylation, and even overall occupancy of the N-glycosylation site at Asn297 was reduced in this mutant form. In fact, there is significant amino acid variation between the naturally occurring human IgG3 allotypes,[180] however, it is still unclear whether allotype-specific N-glycomes of IgG3 differ as well.

#### Experimentally Validating of Glycoimpact

We first examined the highest glycoimpact relation with a high BLOSUM score, Ile/Val; the substitution should have a negligible global structural impact while dramatically changing glycosylation. Consistently, a glycosite-proximal V84I substitution in HIV-gp120 deactivates the virus; otherwise achieved by mutagenic glycosite-ablation.[103] Similarly, the PrP V180I substitution, adjacent to one of two PrP glycosylation sites (N181), is a causal mutation in Creutzfeldt-Jakob disease.[102]

Through multiple evolutionary and pathogenic lenses, we further examined the relevance and veracity of glycoimpact. We found that higher glycoimpact correlates with a divergence between the function-focused PAM and structure-focused BLOSUM substitution matrices. The observation corresponds to a common understanding in glycobiology, that both glycosylation and structure are necessary to explain protein function. Here, we expressed that conclusion as amino acid substitutions and therefore present evidence of a glycan-protein structure association. We further corroborated that glycoimpact explains the divergence between structure and conservation-based predictions by examining dbNSFP pathogenicity scores. Structure and function-oriented pathogenicity tools co-segregate, suggesting glycoimpact is capturing a consistent and definitive difference between these tools. Probing further into evolutionary statistics, we found that over half of the amino acids are significantly coupled with glycosites. These results suggest the N-glycan-determining sequence is larger and more sophisticated than the necessary N-glycan sequon (NX[T/S]).

Observing high-glycoimpact pathogenic mutations close to glycosylation sites is consistent with the bounded biosynthesis hypothesis. Indeed, we found an enrichment of pathogenic mutations close to glycosites in ClinVar. One such glycosite-proximal (site N371) and high-glycoimpact mutation, tyrosinase/A355V (P14679), is sufficient to cause albinism.[52] N371 glycosylation is likely important, because T373K (which ablates N371 glycosylation) is a well-characterized causative mutation in albinism.[53,54,181] Glycosylation of this protein is critical for folding[55,104] and its ablation results in decreased protein abundance and activity.[105] Additionally, non-mutagenic deglycosylation of tyrosinase also interrupts function.[57] In a published structure (PDB:5M8N), A355 is less than 20Å from N371. The literature suggests that disrupting tyrosinase glycosylation is sufficient to induce Albinism. Here, we suggest an additional mutation acts through glycosylation disruption.

#### Demonstration of the plausibility (PrP), accuracy (HIV), specificity (IgG), and reproducibility (SARS-CoV-2) of glycoimpact predictions

In examining human prion protein, HIV envelope, IgG, and SARS-CoV-2 spike glycosylation in the context of glycosite proximal substitutions, we revealed specific instances of functionally important glycoimpact events. Given that glycosylation impacts PrP pathogenicity,[58] the proximity of pathogenic mutations and glycosylation sites in prion disease suggests a mechanism for glycan modulation of pathogenicity. Here we observed that CJD mutations are significantly close to N181 or N197 (7 of 8 occur within 10Å) while GSD mutations only trend closer to this site, suggesting specific genetic and deliberate modulators of these pathogenic glycosylation events. Similarly, common mutations in SARS-CoV-2 spike protein occur close to several glycosylation sites.[71] Through the lens of glycoimpact, glycan-modulated pathogenicity appears more common than originally believed.

Exploring the fidelity of more specific predictions, we compared our predictions to empirical observations of glycan complexity and composition. We successfully predicted glycan complexity measurements on HIV gp160 envelope protein. More specifically, we validated the complexity-modulating effect of two amino acids, P, and Q. While our model predicted increased complexity near lysine, there is evidence to suggest that proximal lysines should promote low-complexity glycans;[182] the incongruity is likely due to the nonlinear effects observed in the HIV g160 data (**Figure** 6c). Towards furthering the specificity of our predictions, we examined specific glycan substructures associated with mouse IgG1 allotypes. We found associations suggesting that upstream (N-terminal) I to F substitution could explain the observed increase in galactosylation while spatial-proximal I to F substitution could explain the observed increase in sialylation.

To mitigate biases related to retrospective data analysis, we produced novel glycoprofiles for Spike proteins across three Variants of Concern (VOC) in SARS-CoV-2. We selected these VOCs due to their high-glycoimpact and glycosite-proximal mutations. As expected, the IMR predicted the differential glycosylation across these three VOCs. If IMR can predict differential glycosylation as viruses evolve, these predictions could be useful in predicting immune evasion and further definitions of the viral evolutionary landscape for many viruses.[14,107,108]

Finally, we examine the functional glycosite-proximal and high glycoimpact variants by examining differential glycosylation and function in Fc variants of well-characterized monoclonal antibodies. We found both fucosylation and fucose-modulated functional response (ADCC) correlated with GEE-IMR core-fucose preference. If IMR can predict differential glycosylation and function in antibodies, we can improve our understanding of Fc-FcR induction of functional immune response.

#### Supplemental Tables

*Supplementary **Table** 1* – Number of ClinVar mutants (source dbNSFP3) by BLOSUM64 (>=0), distance to the nearest glycosite (near<30*Å*), or Glycoimpact (high>2.5).

| Distance to Glycosite | Glyco-Impact | Variants | Proteins | Not Benign | Benign |
|---|---|---|---|---|---|
| Far | High | 195 | 80 | 171 | 24 |
| Far | Low | 357 | 126 | 332 | 25 |
| Near | High | 396 | 179 | 340 | 56 |
| Near | Low | 832 | 285 | 766 | 66 |

*Supplementary **Table** 2* – Prion disease variants in CJD and GSS

| Prion Disease | Variants | Citation |
|---|---|---|
| Creutzfeldt-Jakob disease (CJD) | P105L, Y145X, Y163X, D178N, T188K, E196A, E200K, V201I, M232R | [66,67] |
| Gerstmann-Sträussler-Scheinker disease (GSS) | P102L, A117V, G131V, F198S, Y226X, Q227X | [68,69] |

*Supplementary **Table** 3* – P-values and FDR correction for two-sample Mann-Whitney tests of the glycan abundance distributions (***Figure** 7*c, *Supplementary **Figure** 14*).

| GLYCAN | P-VALUE | FDR |
|---|---|---|
| **G** | 0.000274 | 0.000353 |
| **G1** | 0.007251 | 0.008158 |
| **G2** | 2.04E-06 | 3.06E-06 |
| **S** | 1.02E-06 | 1.84E-06 |
| **S1** | 1.02E-06 | 1.84E-06 |
| **S2** | 1.02E-06 | 1.84E-06 |
| **G0** | 1.02E-06 | 1.84E-06 |
| **B** | 0.436617 | 0.436617 |

*Supplementary **Table** 4* – Distribution and sample size of IMR conditional probabilities *Supplementary **Figure** 7*F

| MOTIF LENGTH | STRUCTURE FIXED | MEAN | STANDARD DEVIATION | N: P(A\|B)>P(A) | N: P(A\|B)<P(A) | N | FOLD CHANGE |
|---|---|---|---|---|---|---|---|
| **1** | glycan | 0.039501 | 0.065099 | 548 | 634 | 1183 | 2.314301 |
| **1** | protein | 0.091417 | 0.13807 | 548 | 634 | 1183 | |
| **2** | glycan | 0.033059 | 0.058625 | 1025 | 1160 | 2185 | 2.519216 |
| **2** | protein | 0.083284 | 0.117493 | 1025 | 1160 | 2185 | |
| **3** | glycan | 0.027672 | 0.054289 | 1576 | 1788 | 3364 | 3.235929 |
| **3** | protein | 0.089546 | 0.115724 | 1576 | 1788 | 3364 | |
| **4** | glycan | 0.025251 | 0.045312 | 1798 | 2056 | 3855 | 3.704429 |
| **4** | protein | 0.093541 | 0.120251 | 1798 | 2056 | 3855 | |
| **5** | glycan | 0.025405 | 0.04588 | 2327 | 2628 | 4956 | 3.470337 |
| **5** | protein | 0.088163 | 0.104168 | 2327 | 2628 | 4956 | |
| **6** | glycan | 0.024508 | 0.047592 | 2941 | 3369 | 6311 | 3.562296 |
| **6** | protein | 0.087305 | 0.098485 | 2941 | 3369 | 6311 | |
| **7** | glycan | 0.023376 | 0.045586 | 4131 | 4761 | 8893 | 4.097588 |
| **7** | protein | 0.095787 | 0.103835 | 4131 | 4761 | 8893 | |
| **8** | glycan | 0.022285 | 0.041808 | 4898 | 5711 | 10612 | 4.714163 |
| **8** | protein | 0.105054 | 0.109607 | 4898 | 5711 | 10612 | |
| **9** | glycan | 0.021484 | 0.037508 | 4913 | 5742 | 10658 | 5.043449 |

| | | | | | | | |
|---|---|---|---|---|---|---|---|
| **9** | protein | 0.108351 | 0.11061 | 4913 | 5742 | 10658 | |
| **10** | glycan | 0.020017 | 0.032146 | 4319 | 5183 | 9506 | 5.641682 |
| **10** | protein | 0.11293 | 0.113366 | 4319 | 5183 | 9506 | |
| **11** | glycan | 0.019036 | 0.026794 | 3525 | 4367 | 7895 | 5.758148 |
| **11** | protein | 0.109612 | 0.107277 | 3525 | 4367 | 7895 | |
| **12** | glycan | 0.017643 | 0.022947 | 2579 | 3304 | 5885 | 5.907276 |
| **12** | protein | 0.104222 | 0.096894 | 2579 | 3304 | 5885 | |
| **13** | glycan | 0.015433 | 0.018858 | 1894 | 2475 | 4371 | 6.921746 |
| **13** | protein | 0.106823 | 0.096742 | 1894 | 2475 | 4371 | |
| **14** | glycan | 0.012897 | 0.014746 | 1399 | 1856 | 3255 | 8.261073 |
| **14** | protein | 0.106543 | 0.094723 | 1399 | 1856 | 3255 | |
| **15** | glycan | 0.010178 | 0.011166 | 931 | 1253 | 2185 | 11.10655 |
| **15** | protein | 0.113045 | 0.097098 | 931 | 1253 | 2185 | |
| **16** | glycan | 0.008308 | 0.008946 | 623 | 835 | 1458 | 14.74224 |
| **16** | protein | 0.122482 | 0.102175 | 623 | 835 | 1458 | |
| **17** | glycan | 0.006909 | 0.007713 | 427 | 565 | 993 | 19.85786 |
| **17** | protein | 0.137195 | 0.108164 | 427 | 565 | 993 | |
| **18** | glycan | 0.006178 | 0.006757 | 275 | 359 | 634 | 22.56627 |
| **18** | protein | 0.139421 | 0.107794 | 275 | 359 | 634 | |
| **19** | glycan | 0.004446 | 0.004865 | 158 | 214 | 372 | 31.24682 |
| **19** | protein | 0.13892 | 0.107965 | 158 | 214 | 372 | |
| **20** | glycan | 0.004446 | 0.004865 | 158 | 214 | 372 | 31.24682 |
| **20** | protein | 0.13892 | 0.107965 | 158 | 214 | 372 | |
| **21** | glycan | 0.00363 | 0.003766 | 118 | 161 | 279 | 34.24255 |
| **21** | protein | 0.124301 | 0.101235 | 118 | 161 | 279 | |

235

#### Supplemental Figures

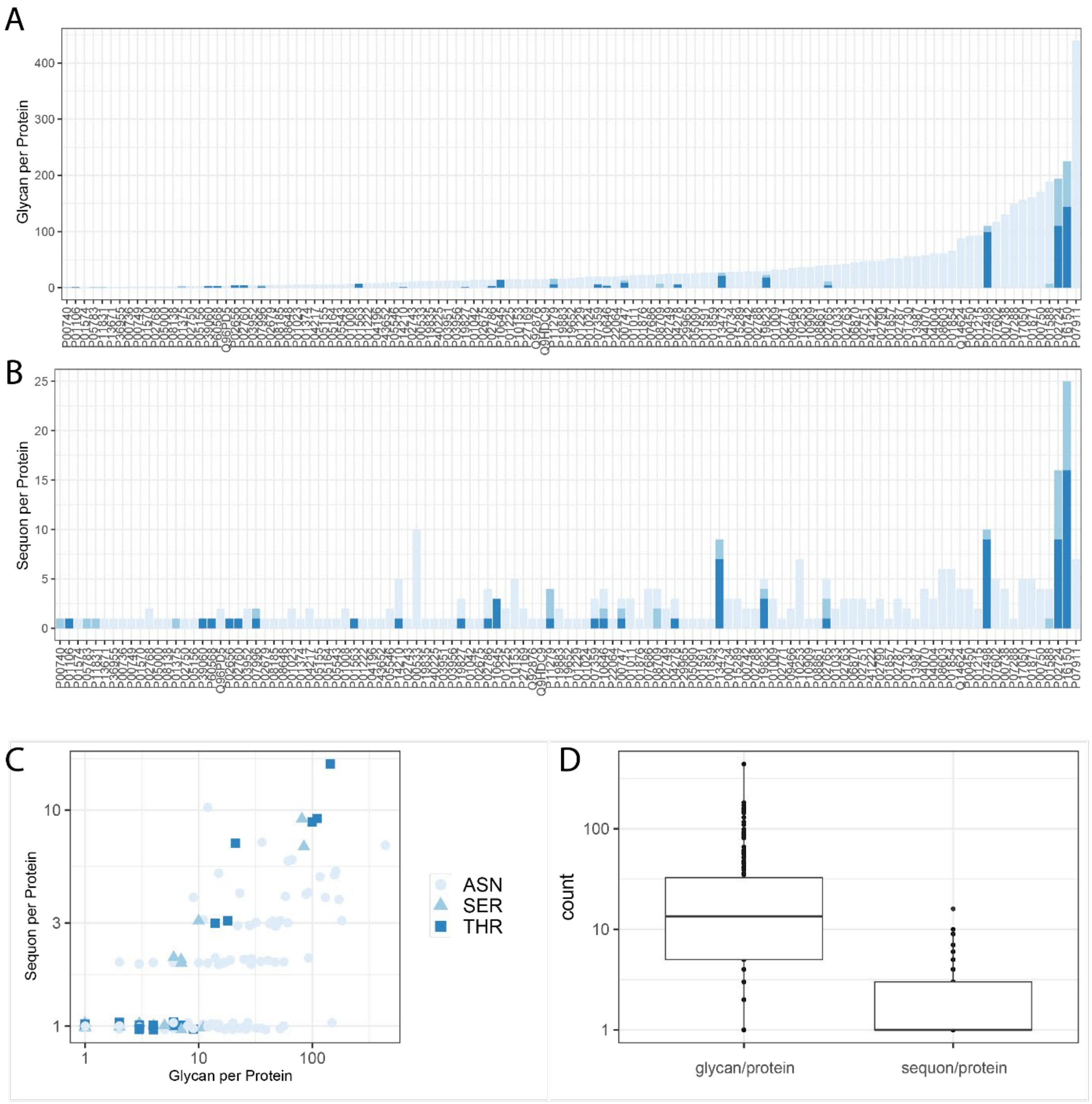


Supplementary **Figure** 1 – PGD proteins, glycans, and sequons. (**A**) Number of glycans assigned to each protein in PGD. (**B**) Number of sequons described within each protein. (**C**) Pairwise comparison for each protein of number of sequons and glycans assigned to each protein. (**D**) Direct comparison of the glycan and sequon per protein in PGD.

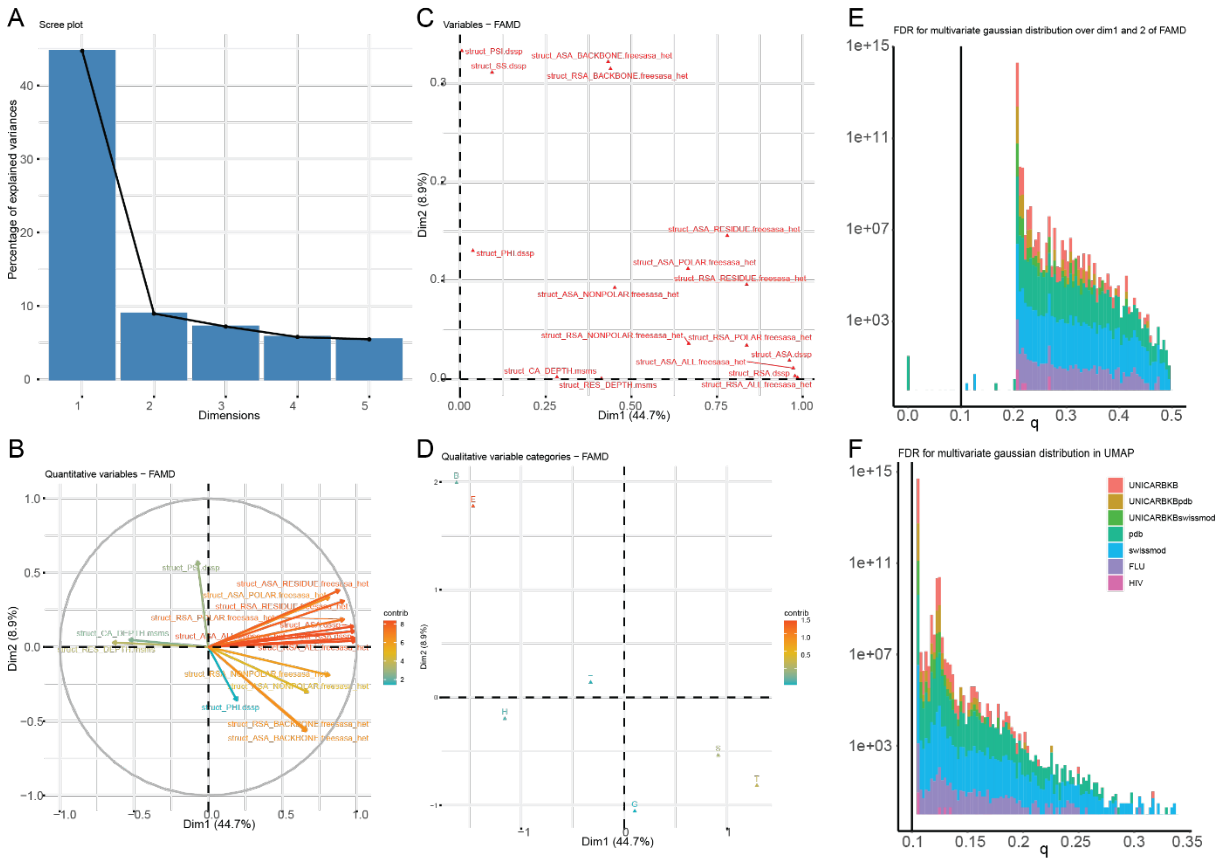


*Supplementary* ***Figure 2*** – Site-specific structure and chemistry diversity among glycoproteins examined. (**A**) Screen map depicting the variance represented by each dimension of the Factor Analysis with Mixed Data (FAMD).[183] (**B-D**) Feature weights from a FAMD of annotated sites. (**C**) FDR distribution of p-values from a multivariate gaussian of the (**E**) FAMD (first two dimensions) and UMAP 2D projection.

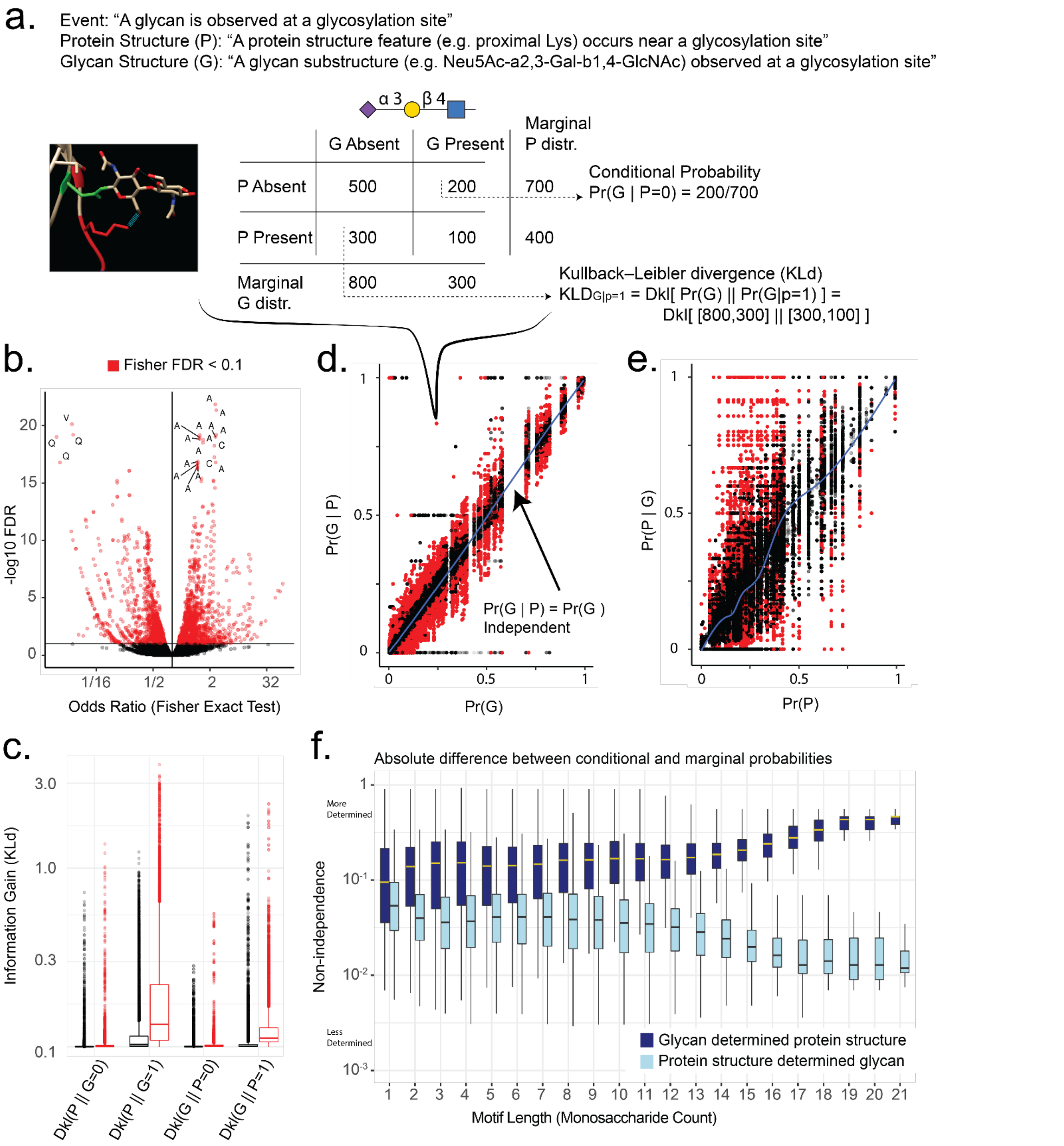


*Supplementary* ***Figure*** *3* – **N- and O-glycan substructures associated with glycosite-proximal protein structure.** (**a**) Specification of the event space to each observed glycosylation event at a glycosylation site. (**b**) Volcano plot of the log odds ratio and False Discovery Rate adjusted p-values from a Fisher exact test between glycan and protein structure occurrence. We observe many large and significant Intramolecular Relations (IMR). (**c**) Kullback-Leibler divergence when either glycan structures, G, or protein structures, P, are specified as present, 1, or absent, 0. For significant IMR (Fisher exact test, FDR<0.1, red), the presence of G or P substantially decreases uncertainty in the unknown variable. (**d**) Probabilities of glycan structures when protein structures were known or “fixed.” **(e)** Protein structure probabilities conditioned on fixed glycan structures. As expected, significantly non-independent IMR (Fisher exact test, FDR<0.1, red) fall consistently off the diagonal**. (f)** Non-independence (absolute difference between conditional and marginal probabilities |Pr(A|B)-Pr(A)|) stratified by glycan motif size for protein-glycan relations when glycan structures are fixed (dark blue; Pr(P|G)-Pr(P)) and when protein structures are fixed (light blue; Pr(G|P)-Pr(G)). This panel only includes IMR significant in the Fisher test.

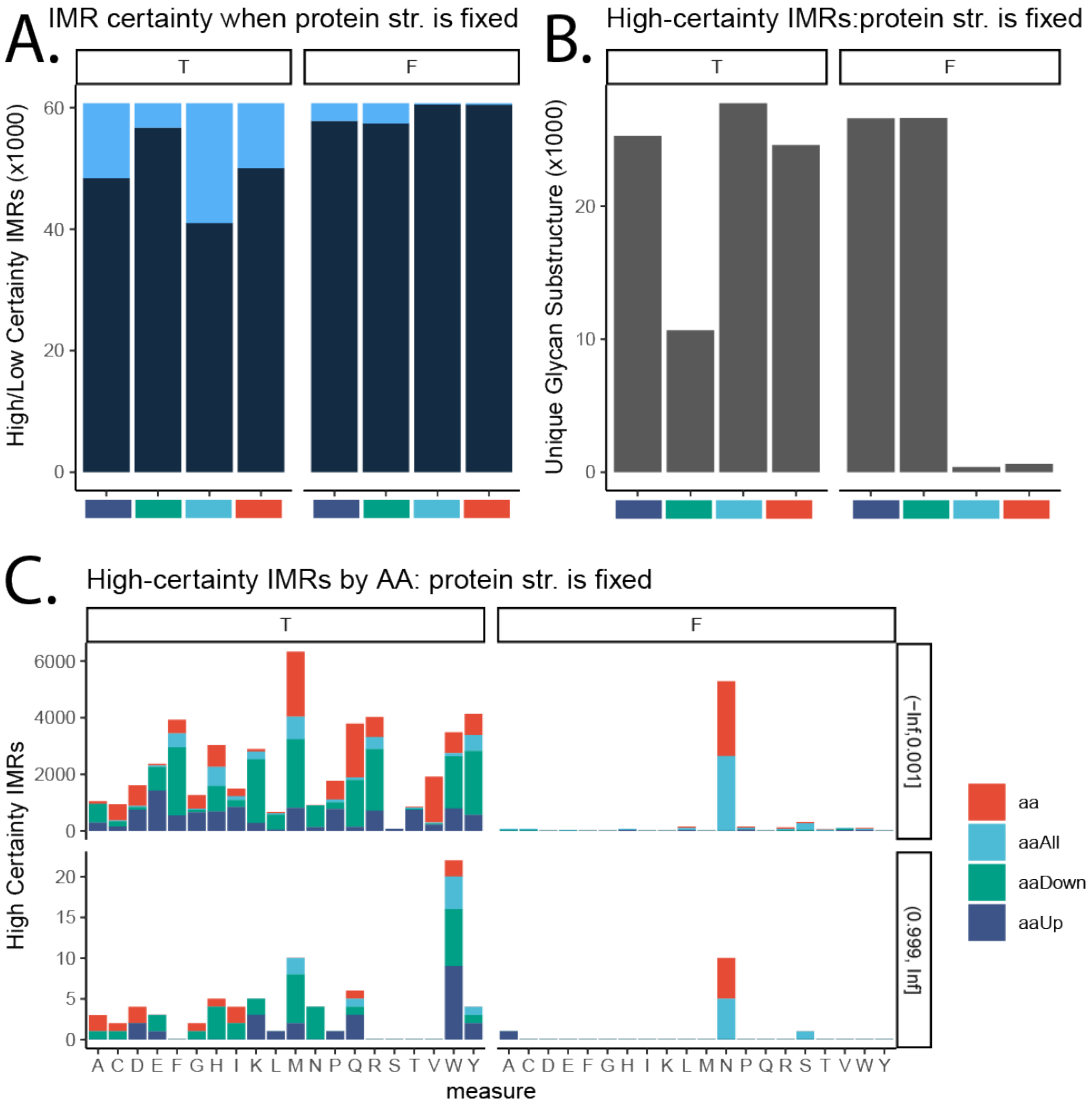


*Supplementary* ***Figure*** *4* - Distributions of high-confidence (close to 1 or 0; within 0.001) amino acid-glycan IMR. Each plot is split indicating when the protein structure is fixed present (T, left) or absent (F, right). (**A**) Division of high-confidence (light blue) and other (dark blue) amino acid -glycan IMR. (**B**) Number of unique glycan substructures involved in high-confidence amino acid - glycan IMR. (**C**) High-certainty aa-glycan IMR by amino-acid (x-axis) proximity type (color) and probability: close to zero (top) or close to one (bottom).

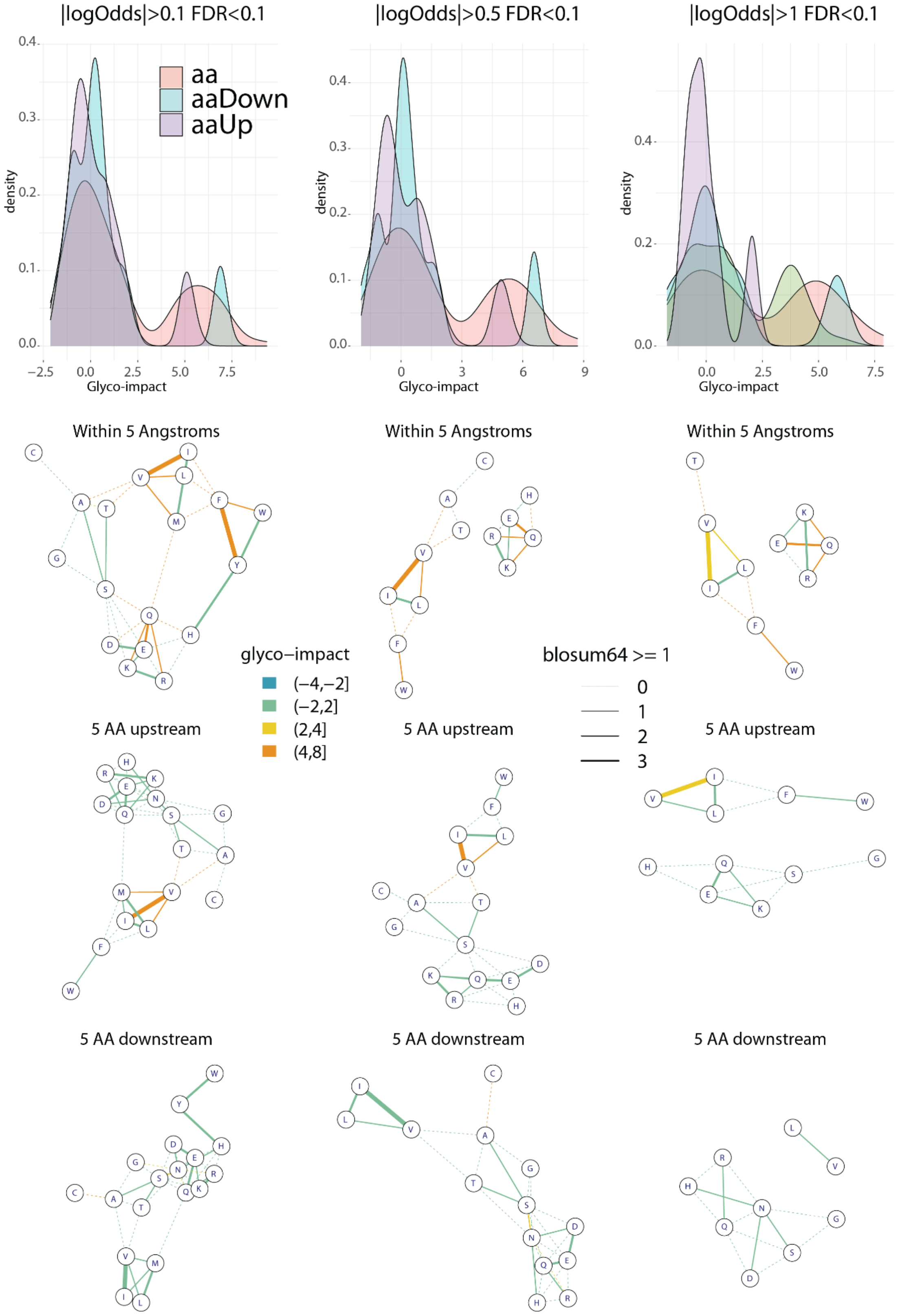


*Supplementary **Figure** 5* – Glycoimpact at various IMR thresholds.

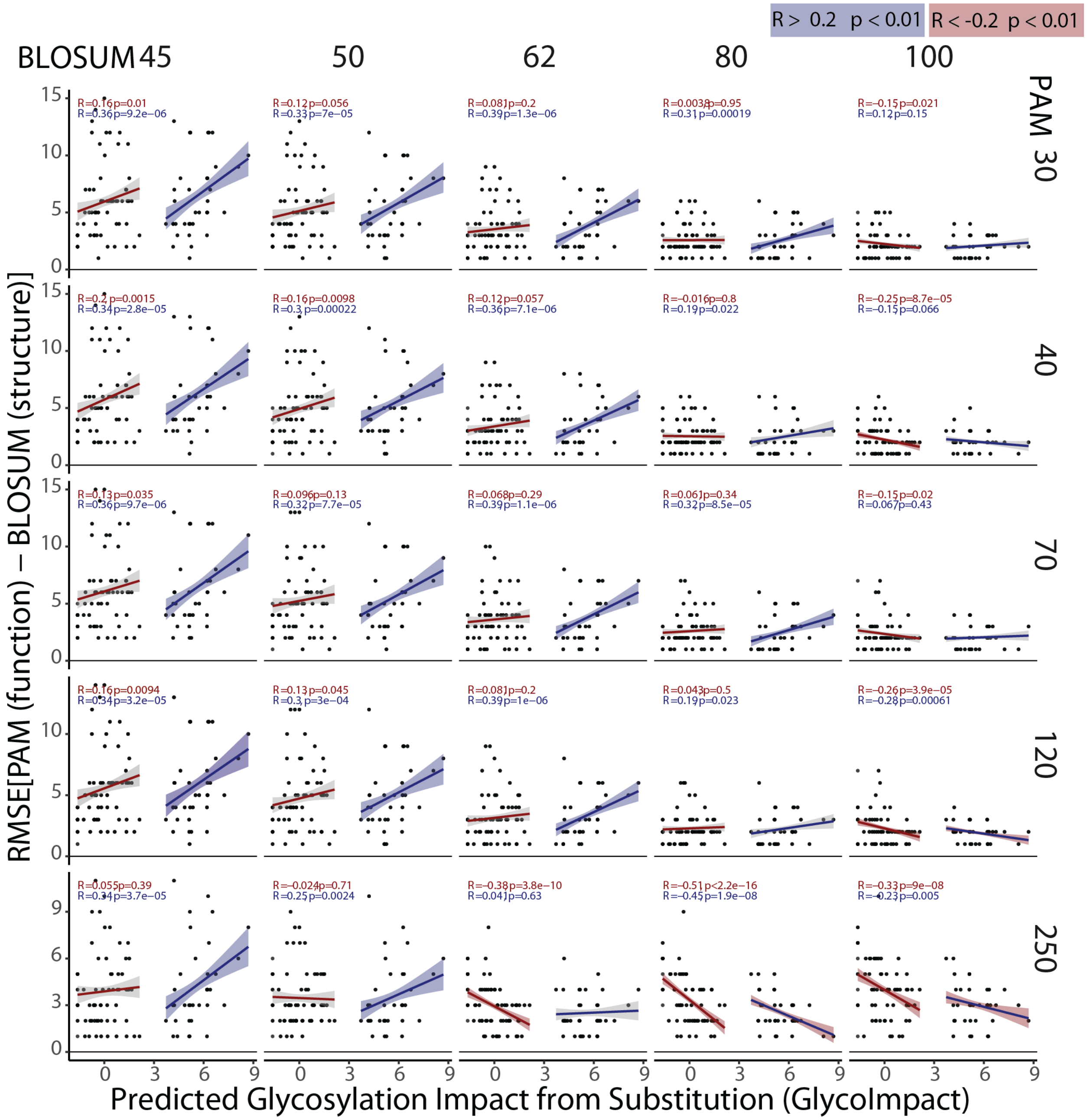


*Supplementary **Figure** 6* – Root mean square error between PAM and BLOSUM substitution scores correlated with predicted glycoimpact.

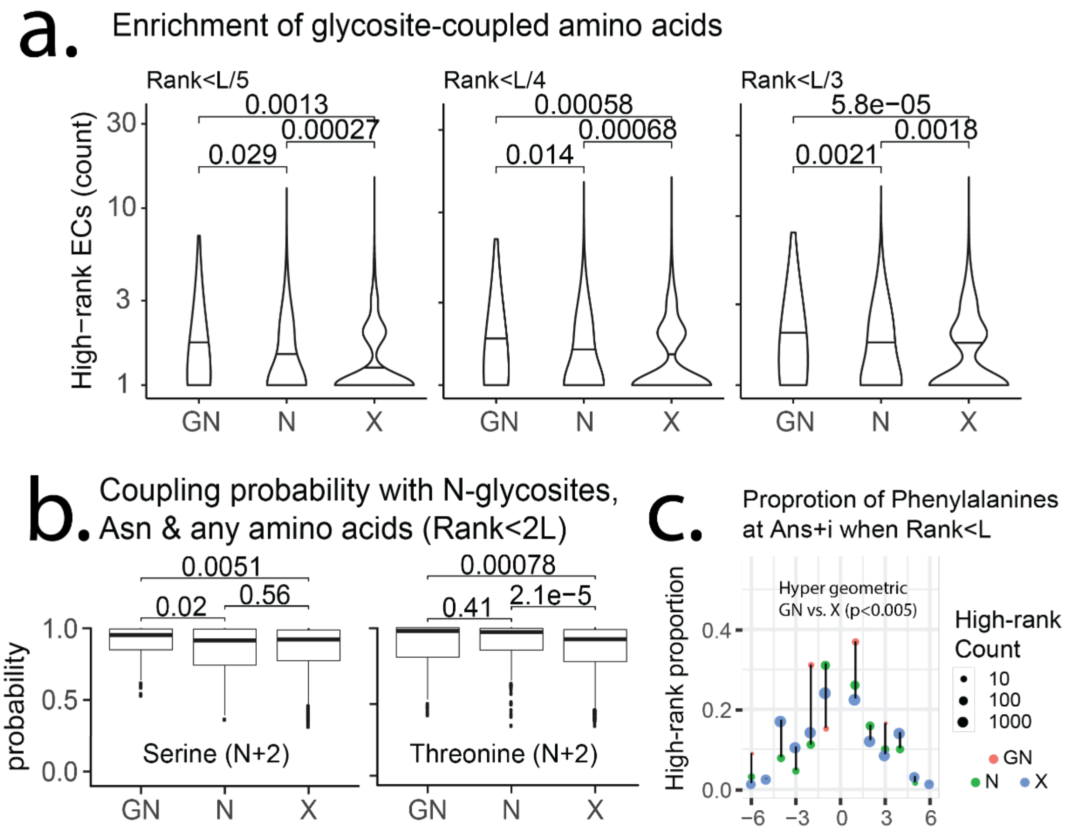


*Supplementary* ***Figure*** *7 – Evolutionary coupling with glycosites.* (**a**) The number of high-ranking (p<L/5-3) evolutionary coupling events between glycosites (GN), asparagines (N), or any residue (X) with all other residues in each of 2,005 alignments. (**b**) Evolutionary coupling (EC) probability between serine, threonine or histidine with a glycosite (GN), any asparagine (N), or any amino acid (X). Serines and threonines considered appear two residues C-terminal to the GN, N or X (N+2). P-values were calculated using a one-sided Wilcoxon test. (**c**) Illustrating hypergeometric enrichment at each relative position, the proportion of high-ranking ECs (Rank<L) at N+/-i relative to a glycosylation site (GN, pink), asparagine (N, green) or any amino acid (X, blue).

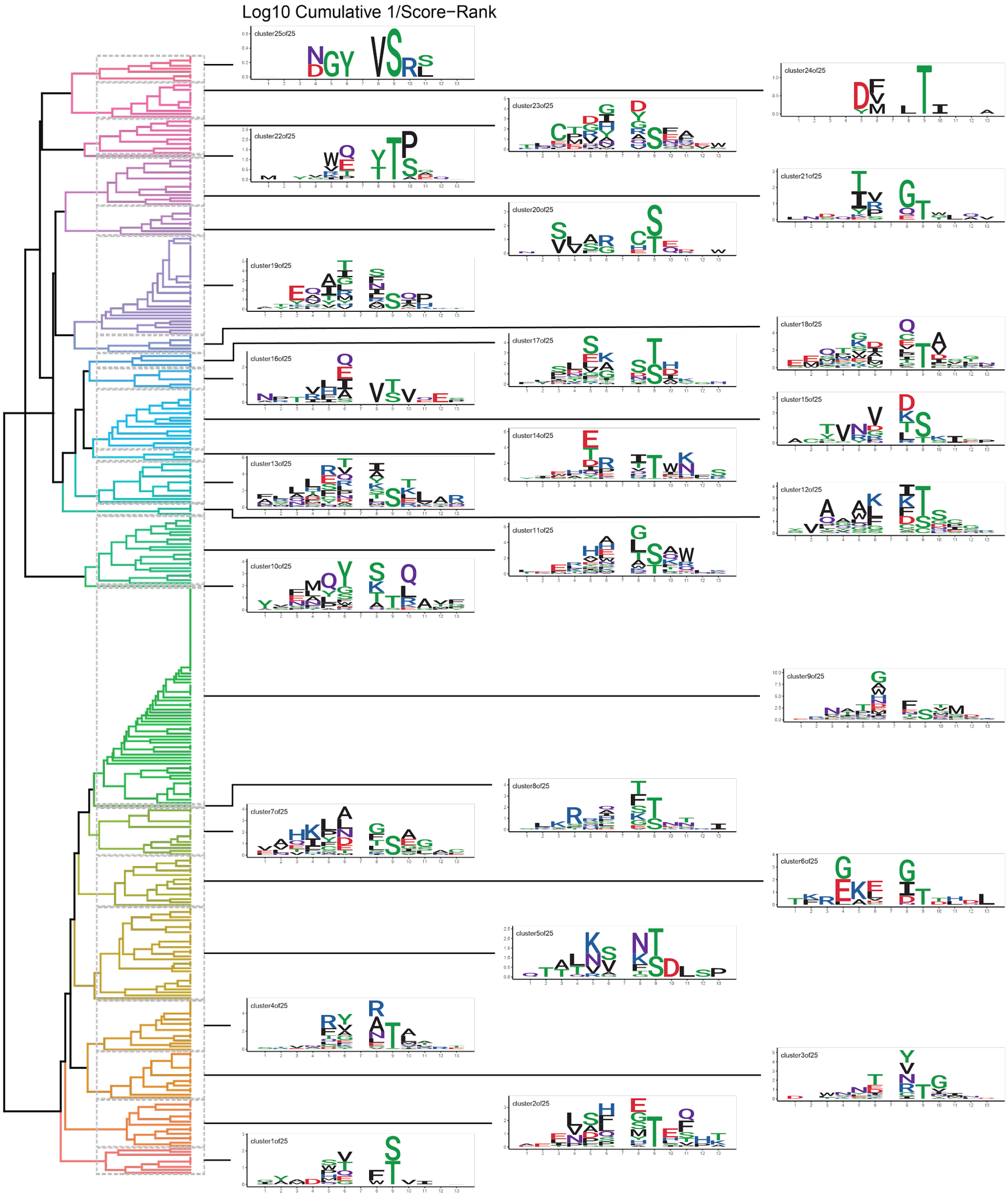


*Supplementary **Figure** 8* - EC-masked extended sequon clustering with motif logos describing high-ranking glycosite-coupled ECs at each position.

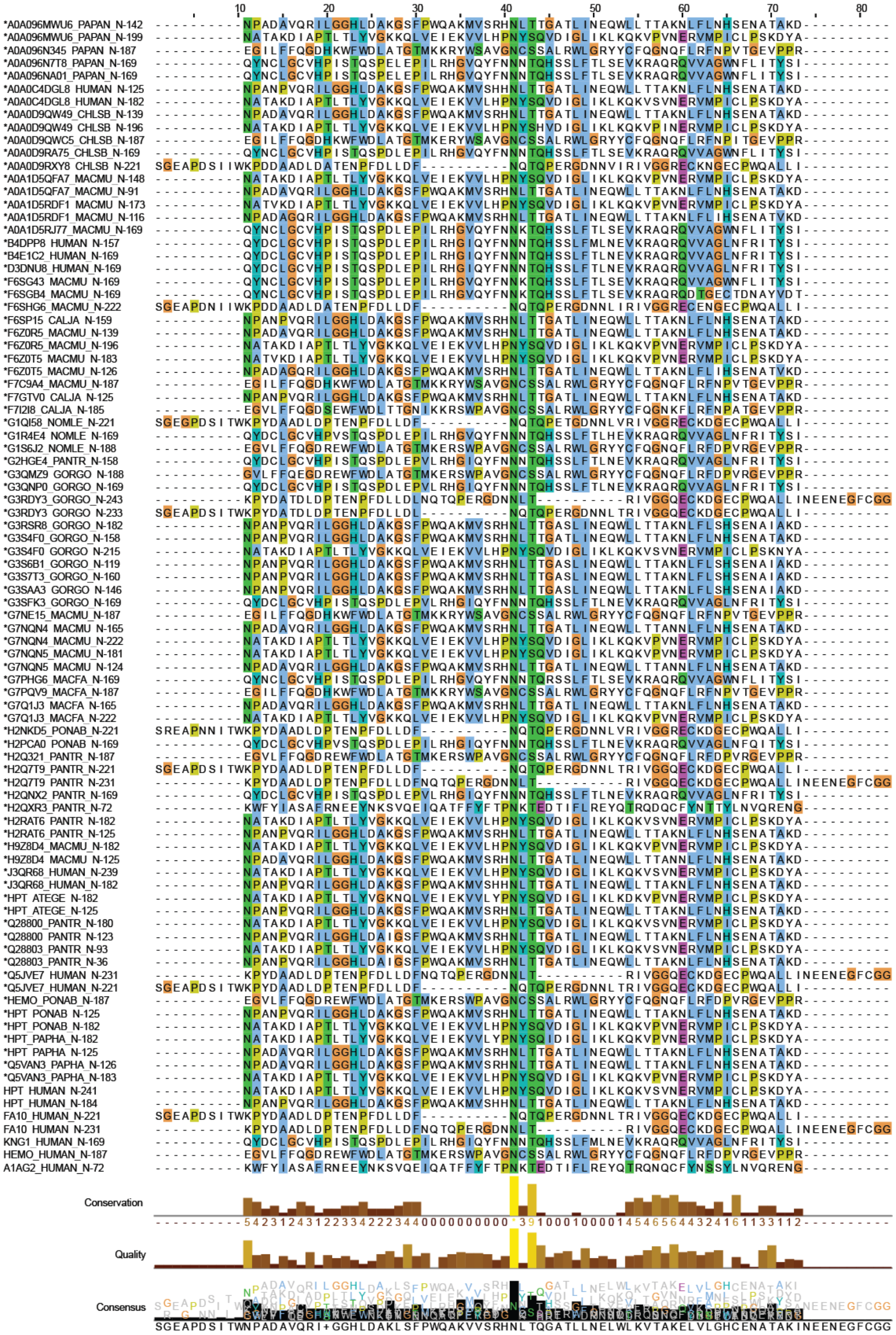


*Supplementary* ***Figure*** *9* - Glycosite alignment for tetraantennary structures with no sialic acid or fucose from GlycoSiteAlign at GlyConnect.

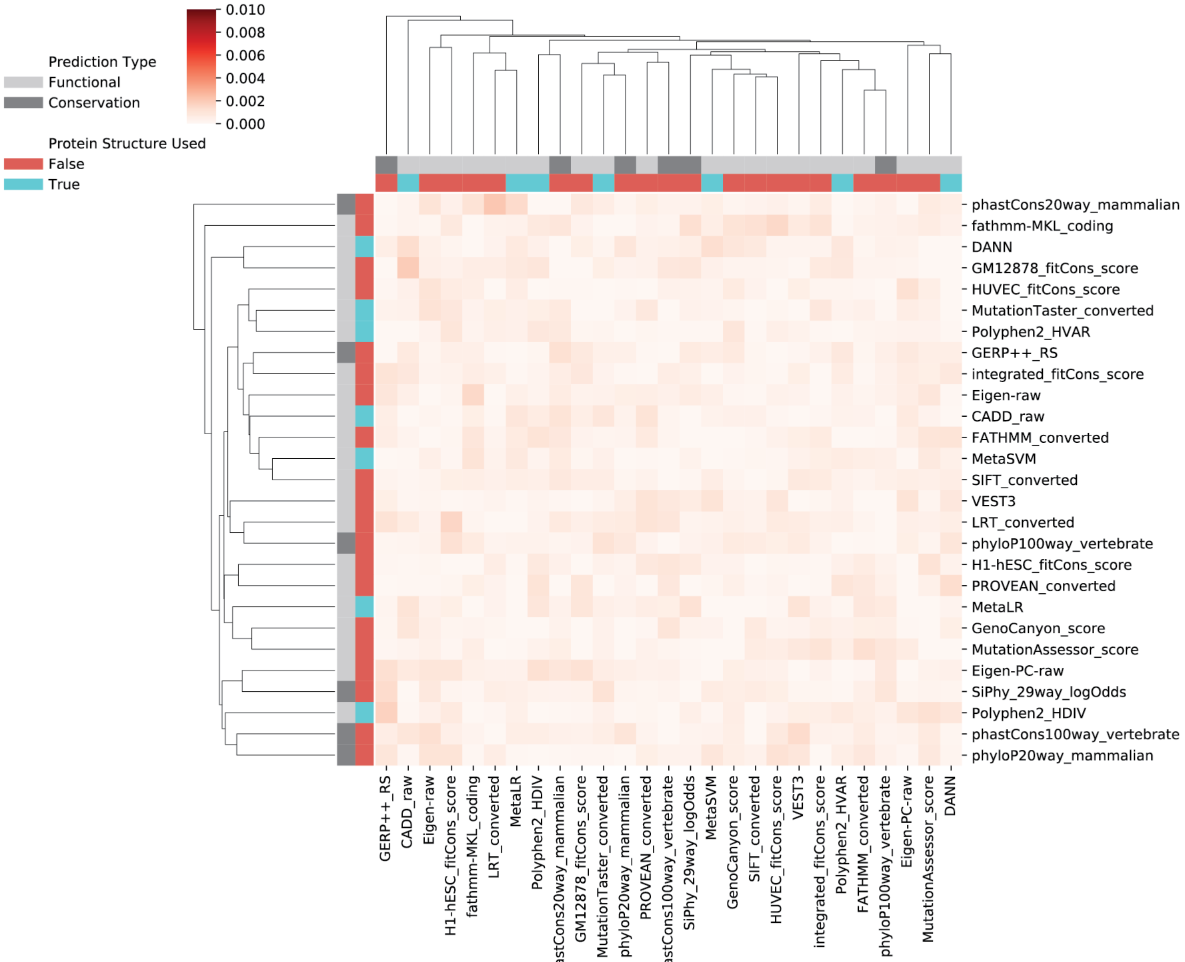


*Supplementary **Figure** 10* – Correlation between glycoimpact and pathogenicity score pairwise errors when pathogenicity scores are shuffled within score.

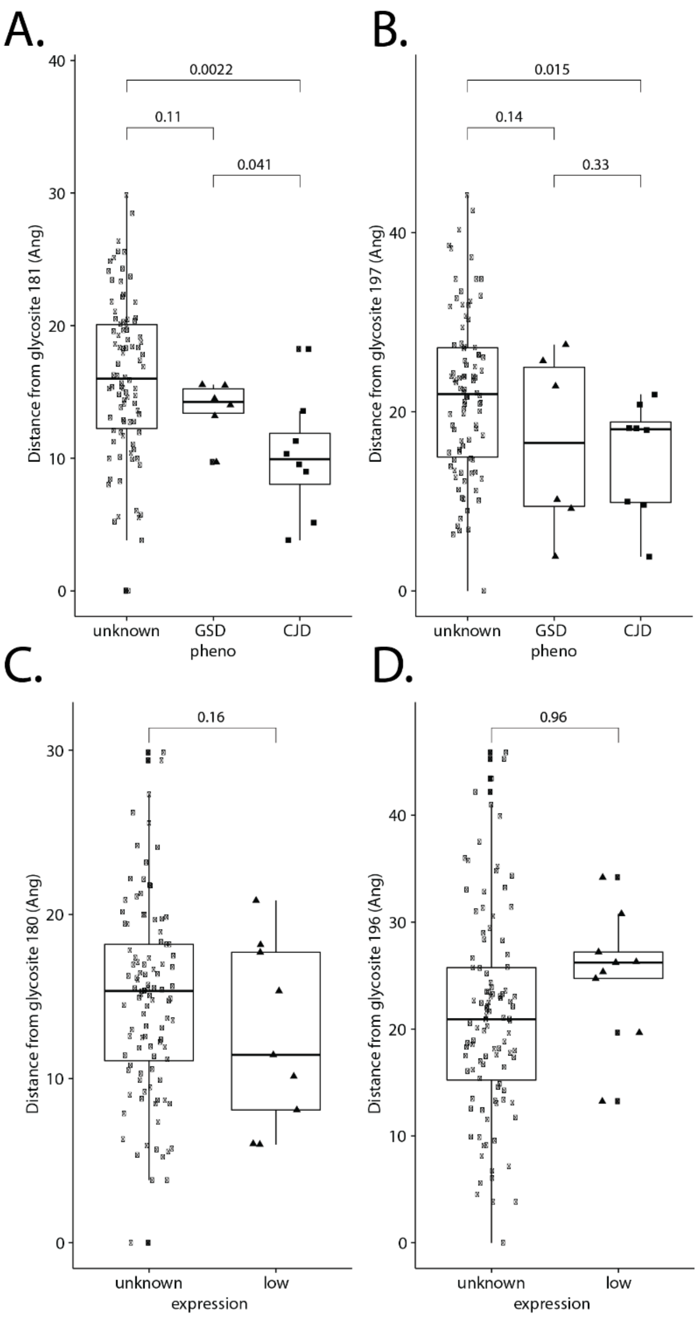


*Supplementary* ***Figure*** *11* – Minimum distance between amino acids within human PrP for disease and library mutants (**A,B**) and low expression mutants (**C,D**).

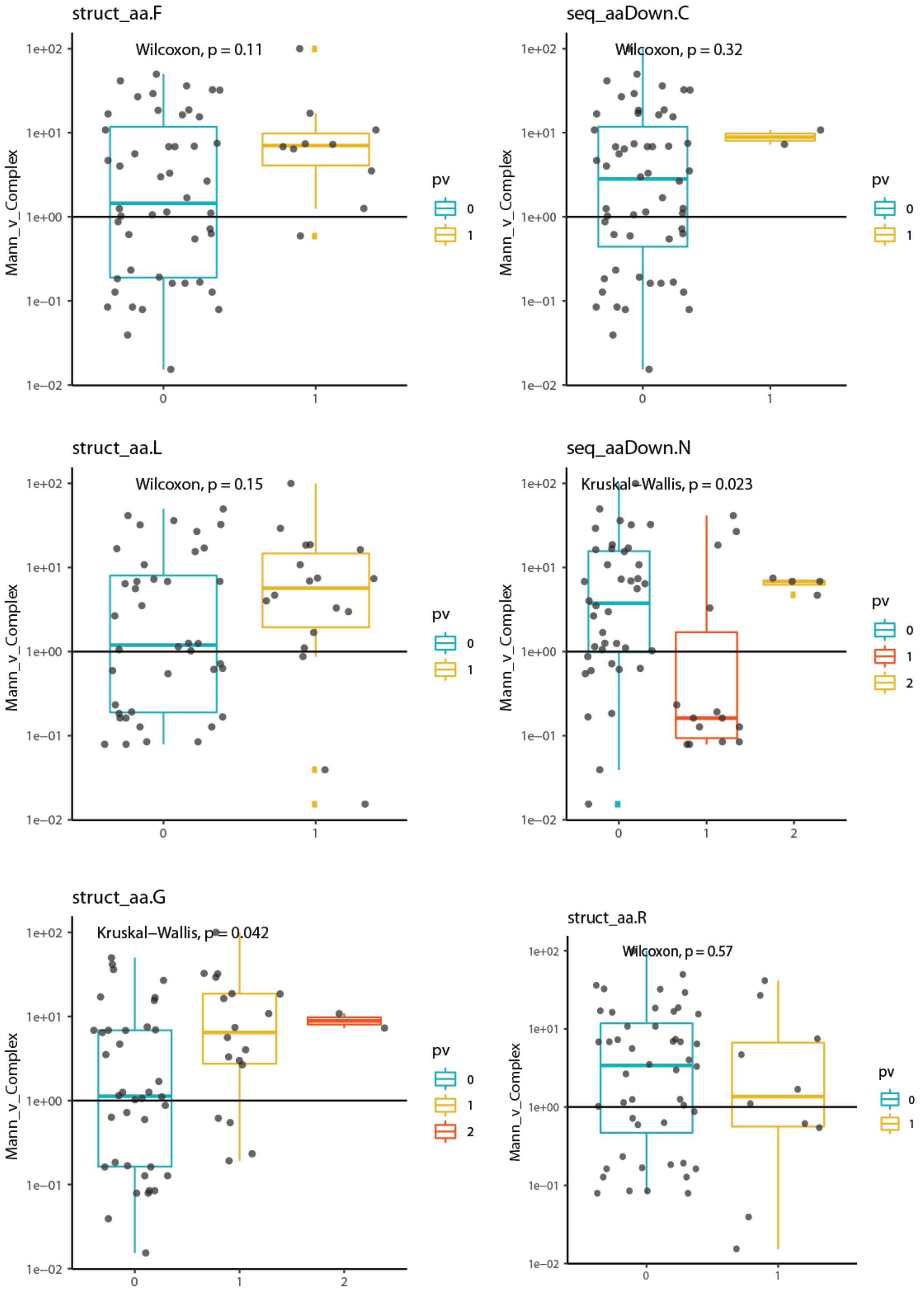


*Supplementary **Figure** 12* – Protein-Glycan structure relations in HIV ENV gp160.

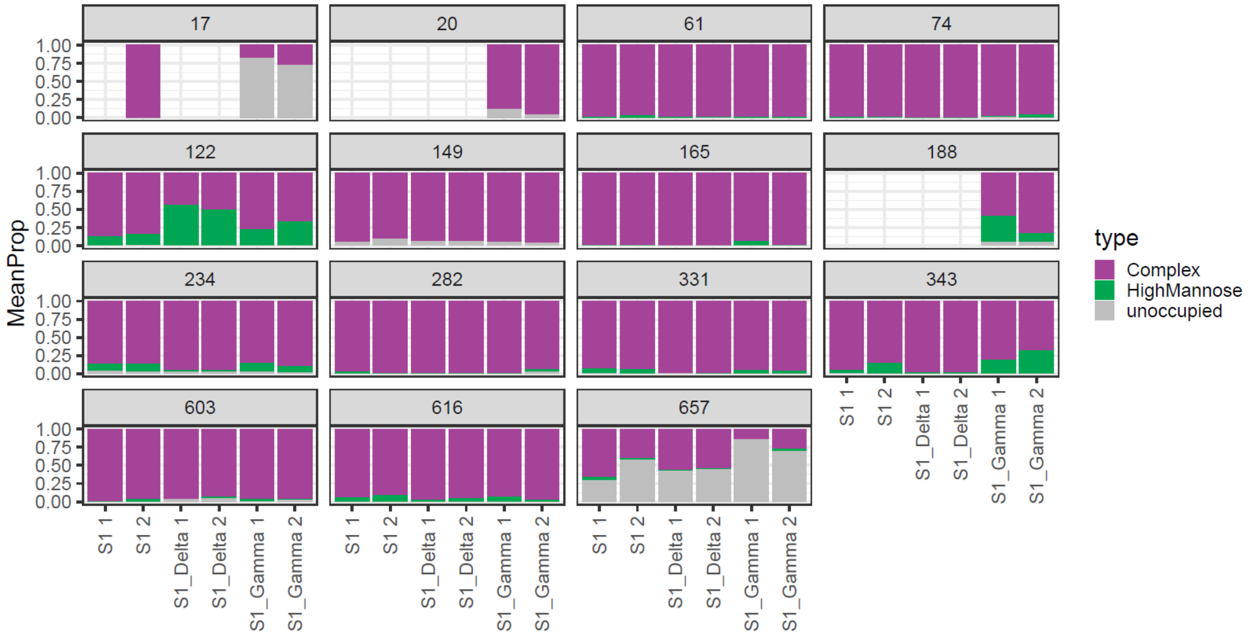


*Supplementary* ***Figure*** *13* - Mean proportion of mass-spectrometry-observed peptides with mass offsets corresponding to Complex, Oligomannose/Hybrid glycans or unoccupied sites in the SARS-CoV-2 S1 subunit across the original 2019 strain, and the Delta and Gamma variants.

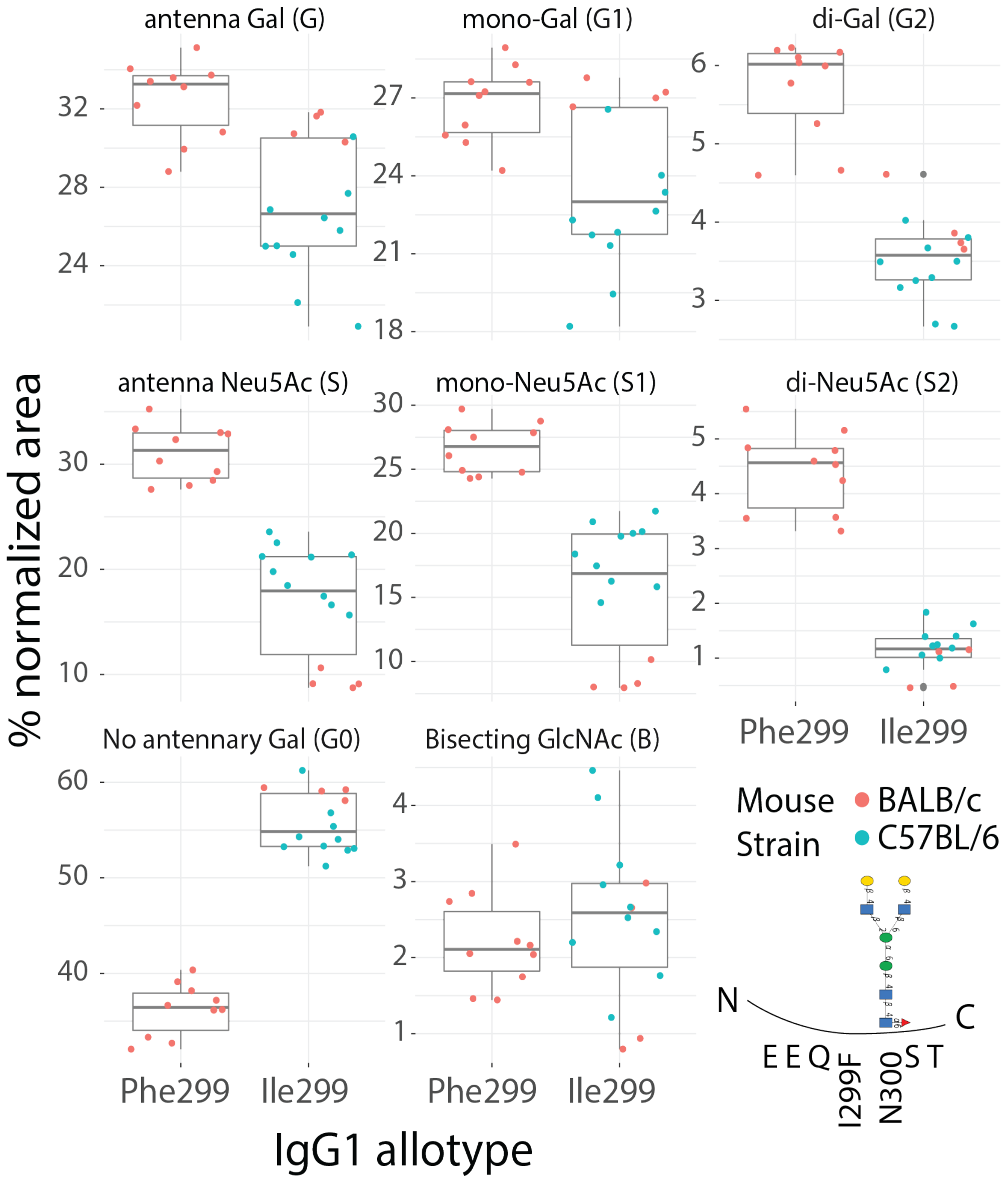


*Supplementary **Figure** 14* – Mouse strain specific IgG3 N-glycosylation measurements; relative abundance.

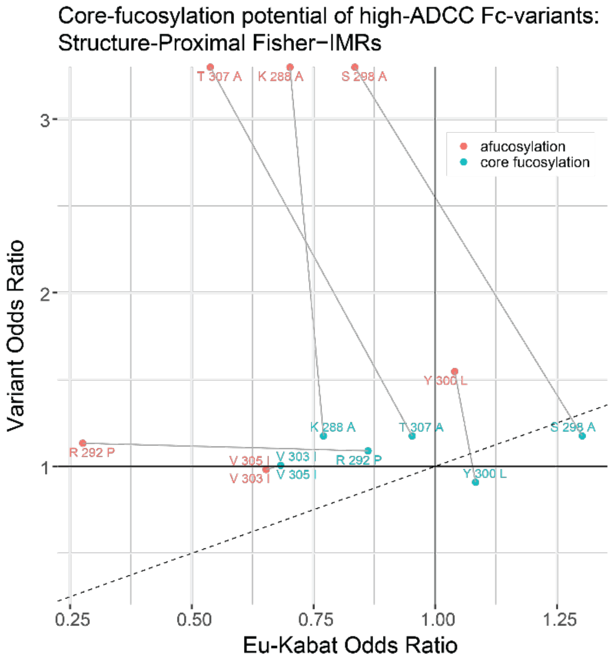


*Supplementary* ***Figure*** *15* - Relative Fisher Odds Ratios (OR) estimated structure-specific IMR, within 6Å, indicating the preference for afucosylation (red) or core-fucosylation (blue) across multiple antibody allotypes. IMR above y=x (line of equality, dotted line) are correlated with the Fc variant and/or anticorrelated with the wild-type. Core-fucosylation preference is the distance from the point representing IMR between the WT and variant amino acids and the afucosylated N-glycan motif (precursor glycomotif, red) to the point representing the same IMR between the WT and variant amino acids and the core-fucosylated N-glycan motif (glycomotif of interest, blue) such that the distance is the component perpendicular to the line of equality.

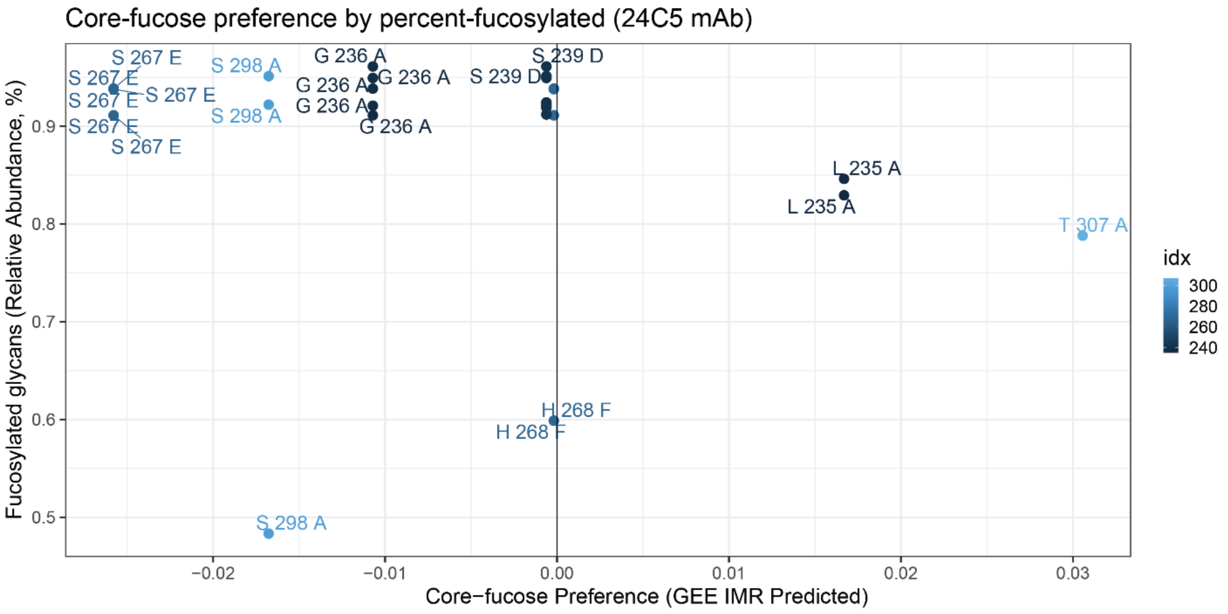


*Supplementary* ***Figure*** *16* – Relationship between IMR-predicted core-fucose preference and relative abundance of fucosylated glycans cleaved from monoclonal antibody 24C5.

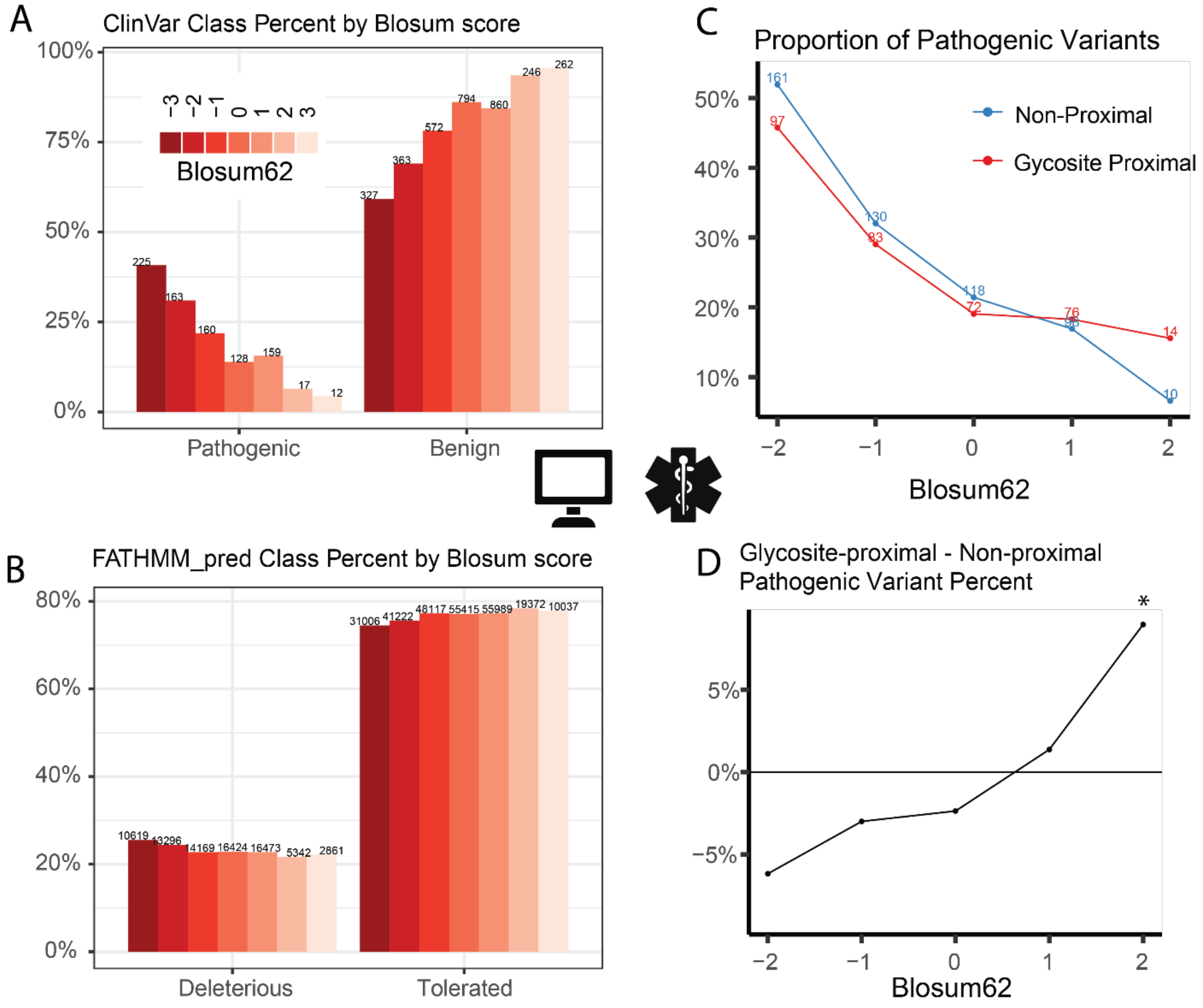


*Supplementary **Figure** 17* - Analysis of ClinVar annotated variants in dbNSFP4.3 stratified by pathogenicity, BLOSUM score and glycosite proximity. (**A**) ClinVar variants stratified by pathogenicity and marginalized by BLOSUM62. (**B**) dbNSFP4.3 variants stratified by FATHMM score and marginalized by BLOSUM62. (**C**) Proportion of pathogenic ClinVar variants marginalized by pathogenicity and BLOSUM score and separated by glycosite proximity. (**D**) Simplified view of panel C showing the difference in proportion between glycosite proximal and non-proximal variants by BLOSUM score. In panel D, Fisher's Exact Test was performed to determine the significance of glycosite proximity on the pathogenic proportion ("*"<0.05).
